# FBH1 Reverses Stalled Replication Forks via Sequential Unwinding of Nascent Strands

**DOI:** 10.64898/2026.06.03.729855

**Authors:** Javier Mendia-García, Emma M. Peacock, Clara Aicart-Ramos, Brandt F. Eichman, Fernando Moreno-Herrero

## Abstract

Replication fork reversal is a DNA damage tolerance mechanism important for genome stability that entails annealing of parental DNA to push the fork backwards. F-box helicase 1 (FBH1) is a 3′–5′ ssDNA translocase and SCF (SKP–CUL1–F box) E3 ubiquitin ligase that catalyzes fork reversal and limits aberrant recombination, yet how its helicase activity drives strand annealing is unknown. Here, using single-molecule and biochemical assays, we show that SCF^FBH1^ reverses forks through a two-stage reaction in which translocation on the lagging strand template while remaining affixed at the junction destabilizes the leading strand duplex to ultimately displace the nascent leading strand. Reversal is force-sensitive and does not generate a four-way junction, revealing an annealing-independent mechanism distinct from those of SMARCAL1, HLTF, and ZRANB3. These results establish the importance of nascent strand unwinding to fork reversal and suggest the existence of distinct pathways that produce unique DNA structures, which has implications for fork restart and its measurement in cells.

## Introduction

Replication forks frequently encounter obstacles that slow or stall DNA synthesis, activating the replication stress response. If unresolved, stalled forks can generate double-strand breaks, making them a potent source of genomic instability(*1*–*3*). Cells employ several tolerance pathways to stabilize and restart stressed forks, including translesion synthesis, repriming, template switching, and fork reversal. In fork reversal, stalled forks are remodeled into a regressed structure resembling a four-way junction by reannealing the parental strands and pairing the nascent strands, which stabilizes the fork and contributes to genome stability(*4*–*6*). Because unprotected or improperly restarted reversed forks are prone to nucleolytic degradation, reversal must be precisely controlled to preserve genome integrity(*7*).

Several enzymes have been shown to catalyze fork reversal *in vivo* and *in vitro*, including SMARCAL1, HLTF, ZRANB3, and FBH1. The biochemical properties of fork reversal have been characterized most extensively for the SNF2-family of double-stranded (ds)DNA translocases, which include SMARCAL1, HLTF, ZRANB3, bacterial RecG, and phage-encoded UvsW(*6, 8*–*16*). These enzymes promote reversal by translocating on the parental duplex while using unique fork recognition domains to provide specificity for the junction and/or unwind the daughter duplexes(*15, 17*–*20*). These enzymes provide the structural and mechanistic framework for how reversal is defined, although it is not clear how they couple strand annealing and unwinding to generate and catalyze branch migration of a four-way junction. In contrast, the DNA helicase FBH1 catalyzes fork reversal through single-stranded translocase activity(*21*–*23*).

FBH1 is a 3′–5′ SF1 helicase of the UvrD family(*24*–*26*) that promotes replication-fork reversal *in vivo* and *in* vitro in an ATPase dependent manner(*22*), making it an archetypal example of helicase-driven fork remodeling. FBH1 is also a component of the SCF (SKP1–CUL1–RBX1) E3 ubiquitin ligase complex (SCF^FBH1^) that ubiquitinates RAD51, which presumably limits the association of RAD51 with chromatin to prevent unscheduled recombination during replication stress(*23, 27*–*34*). FBH1 is recruited to stalled forks by its association with PCNA through its PIP (PCNA-interacting protein) and APIM (AlkB homologue 2 PCNA-interacting) motifs(*35, 36*). Preferential binding of SCF^FBH1^ to ssDNA on the lagging strand template at a fork junction stimulates its helicase activity(*23, 37*) and is required for fork reversal activity(*23*). A cryo-EM structure of SCF^FBH1^ bound to the lagging strand of a fork showed an intimate interaction between FBH1 and the fork junction, disruption of which abrogated fork reversal but not helicase activity(*23*). These data led to a model in which FBH1 reverses forks by remaining bound to the junction while translocating 3′-5′ on the lagging strand template to facilitate parental strand annealing from behind the fork. This would generate a structurally distinct DNA intermediate from the SNF2-family remodelers that anneal parental DNA from in front of the fork lagging strand(*23*), consistent with FBH1-generated reversed forks requiring a unique set of protection factors(*38*) and dependent on RAD54L-mediated branch migration(*39*). Despite this framework, the molecular basis for fork reversal by SCF^FBH1^ and other helicases implicated in fork reversal (e.g., BLM, WRN, RECQ5, FANCM) remains unresolved(*40*–*42*). Specifically, how ssDNA translocation and nascent strand displacement leads to parental strand annealing, and how the structures of the reversed forks differ from those generated by SNF2 remodelers are still unknown.

Here, we define how the helicase activity of SCF^FBH1^ remodels stalled forks and the structural nature of the DNA intermediate it produces. Using single-molecule magnetic tweezers, we show that FBH1 catalyzes fork reversal through a sequential process that begins with lagging strand unwinding and proceeds to leading strand destabilization in a force-dependent manner. Mutational analysis indicates that interaction of FBH1 with the junction branchpoint is required for the transition into the leading strand unwinding step. Ensemble biochemical studies show that the presence of a 3′ ssDNA overhang on the leading strand template is sufficient for fork reversal of model substrates *in vitro*, and that the presence of ssDNA on both the lagging strand template and the nascent leading strand leads to increased fork reversal activity. We also observe the presence of two FBH1 molecules bound to a single DNA junction, consistent with unwinding both nascent strands. Importantly, AFM imaging and biochemical assays establish that FBH1 does not generate a canonical four-way junction, indicating that its product is fundamentally distinct from those formed by SNF2-family translocases.

## Results

### SCF^FBH1^ catalyzes reversal by unwinding both strands of the fork

We directly visualized SCF^FBH1^-catalyzed fork reversal by using magnetic tweezers to monitor, in real time, the extension of individual DNA molecules that mimic stalled replication forks. In this assay, changes in molecular extension reflect conformational transitions; canonical fork reversal of this substrate leads to a measurable decrease in molecular extension during formation and migration of a four-way junction (**Fig. 1A**). The DNA construct consists of two identical dsDNA arms arranged in an inverted orientation and joined by a short hairpin, generating a three-way junction that permits branch migration (**Fig. 1A, Fig. S1**). To enable engagement by SCF^FBH1^, the substrate contains an 8-nt ssDNA gap on the nascent lagging strand, a feature required for SCF^FBH1^-driven reversal(*23*). The applied force opposes fork reversal and favors restoration of the fork. This assay was previously validated by detecting robust reversal by HLTF and RecG on an ungapped version of the substrate(*23*). We confirmed that HLTF also reverses the gapped substrate (**Fig. S1**).

**Fig. 1.**
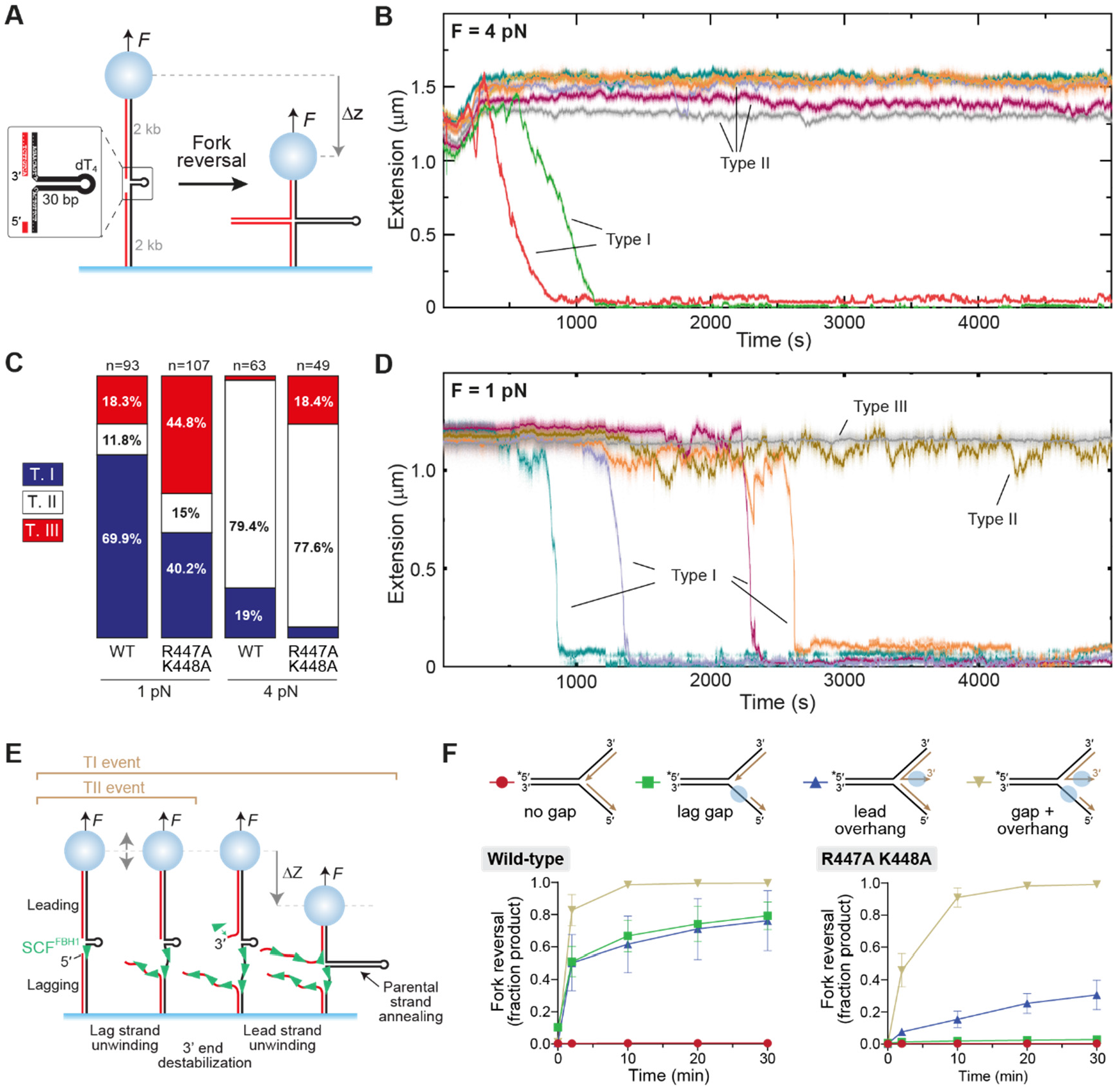
SCF^FBH1^ processively reverses a replication fork-like substrate containing an 8 nt ssDNA gap in the lagging strand. **A**. Schematic of the magnetic tweezers assay. Parental (template) and nascent strands are represented by black and red lines, respectively. Fork reversal is detected as a measurable decrease in DNA extension. *Inset*: detailed view of the three-way junction design for SCF^FBH1^experiments, which contains a 30-bp stem and an 8-nt ssDNA gap in the lagging strand. **B**. Representative time courses of SCF^FBH1^ activity under 4 pN opposing force, showing typical Type I and Type II events (see main text). **C**. Distribution of event types observed for SCF^FBH1^ wild type and R447A K448A mutant at 1 pN and 4 pN forces (number of events is indicated over each bar). **D**. Representative time courses showing individual SCF^FBH1^ activities on the replication fork-like substrate at 1 pN force, illustrating three distinct behaviors (Type I–III events). **E**. Schematic illustrating data interpretation. Sequential unwinding of both leading and lagging strands leads to annealing of the parental strands (black). **F**. Fork reversal by SCF^FBH1^ is stimulated by binding to 3′-ssDNA on the nascent leading strand. Quantification of fork reversal activity of SCF^FBH1^ wild-type (left plot) or R447A K448A mutant (right plot) on the four substrates shown at the top. Data points (mean ± SD, n=4 for wild-type and n=3 for R447A K448A) are quantified from native PAGE separation of substrate and products (**Fig. S5**).

We first monitored fork reversal under an opposing force of 4 pN, which disfavors reversal but still allows branch migration (**Fig. 1B**). Reactions were initiated by introducing SCF^FBH1^ and ATP, and individual tethers were tracked for 90 min. Single-molecule trajectories fell into three reproducible classes, which we refer to as Types I-III (**Fig. 1B-C, Fig. S2**). Type I events (∼19%) displayed a biphasic profile consisting of an initial increase in extension followed by a continuous decrease to the surface, consistent with fork reversal (**Fig. S2**). Type II events (∼79%) showed only the initial extension increase without subsequent shortening (**Fig. S2**), and Type III events (2%) exhibited no detectable change in extension.

The initial increase in extension is indicative of ssDNA generation, which has a greater extension per nucleotide than dsDNA under these conditions (high protein concentration). The magnitude of the extension increase closely matched that measured in an independent unwinding assay performed at the same force (0.12 ± 0.01 nm/nt *vs*. 0.15 ± 0.02 nm/nt (n=33); **Fig. S3**), supporting the interpretation that this phase reflects DNA unwinding accompanied by binding and translocation of SCF^FBH1^ on the generated ssDNA. Given the substrate design, this step most likely corresponds to unwinding of the lagging strand arm. In Type I events, the subsequent decrease in extension reflects entry into a second activity that shortens the tether, which we interpret as leading strand unwinding and subsequent parental strand annealing. Type II events likely correspond to molecules in which lagging strand unwinding occurs, but the reaction fails to progress to this second step.

Since FBH1 does not show any annealing activity at high forces (15 pN)(*23*) and moderate fork reversal at 4 pN, we determined how reversal activity is affected by force by repeating the experiments at 1 pN. Under these conditions, trajectories again grouped into three classes, but their characteristics differed markedly (**Fig. 1C-D, Fig. S4**). Type I events now represented most molecules (∼70%) and resulted in complete shortening to the surface, consistent with fork reversal. These events were typically preceded by a phase of small extension fluctuations without net shortening (86% of type I events). Type II events (∼12%) exhibited similar fluctuations but did not progress to shortening, while Type III events (∼18%) remained inactive. At this lower force, sustained increases of extension were not observed. Instead, transient excursions above baseline occurred in ∼60% of Type I and II events (**Fig. S4**). Independent unwinding assays reproduced these transient increases and oscillations followed by collapse, consistent with cycles of ssDNA generation and secondary structure formation (**Fig. S3**). These observations support the interpretation that the initial fluctuating phase reflects ssDNA generation, although its signal is partially masked at low force. Comparison of event distributions across forces revealed a strong dependence of reaction outcome on mechanical load (**Fig. 1C**). Reducing the force from 4 pN to 1 pN increased the fraction of Type I (reversal) events from ∼20% to ∼70% and reduced Type II events from ∼80% to ∼12%. These shifts indicate that higher force selectively impairs progression from lagging strand unwinding to the second step.

A key distinction emerged when comparing SCF^FBH1^ with HLTF and RecG, both of which produce a four-way junction that is energetically equivalent to the starting fork, allowing reversible branch migration(*23*) (**Fig. S1**). In contrast, SCF^FBH1^-mediated reversal was irreversible under identical conditions, indicating that it generates a structurally distinct product. The model that emerges from the magnetic tweezers data is that FBH1-catalyzed fork reversal proceeds through an initial unwinding of the nascent strands, followed by rapid and spontaneous annealing of parental DNA (**Fig. 1E**), rather than through concerted branch migration of a four-way junction.

This model predicts that progression to the shortening (annealing) phase requires access to ssDNA on the nascent leading strand. If this step is limiting, then providing a pre-existing ssDNA region on the nascent leading strand should facilitate fork reversal. To test this, we used an ensemble assay to compare fork reversal activity of SCF^FBH1^ on forks containing either no gap, a ssDNA gap on the lagging strand template (*lag gap*), a 3′-ssDNA overhang on the nascent leading strand (*lead overhang*), or both (*gap+overhang*) (**Fig. 1F, Fig. S5**). As previously observed, the protein had no activity on the *no gap* fork(*23*). We found that the protein had the same activity for the *lag gap* and *lead overhang* substrates, and that activity was increased when both ssDNA binding sites were present (**Fig. 1F**). This result confirms that unwinding the nascent leading strand facilitates fork reversal by SCF^FBH1^ and is consistent with the shortening phase observed in the magnetic tweezers assay corresponding to leading strand unwinding.

To further define the molecular requirements for FBH1-mediated fork reversal, we examined a mutant (R447A K448A) previously shown to exhibit a severe defect in fork reversal despite retaining helicase activity(*23*). R447 and K448 lie within a positively charged surface patch that contacts the branchpoint of the fork junction in the wake of the motor, indicating that the mutant disrupts engagement with the fork structure rather than compromising core motor function. At 4 pN, the defect was more pronounced, with Type I events reduced to ∼5% while the frequency of Type II was maintained (**Fig. 1C**). The delay before detectable activity (t_0_) was also greatly increased at this load (**Fig. S6**), indicating impaired initiation of productive engagement with the fork. At 1 pN, the defect was less pronounced: the mutant still showed a reduction in Type I events (∼40% vs. ∼70% for wild type), while Type II frequencies remained similar (**Fig. 1C**). Extension increases above the initial value were still observed, indicating that lagging-strand unwinding remains intact (**Fig. S4, Fig. S6**). Kinetic parameters were nonetheless altered, with the mutant displaying longer delays before detectable activity (t_0_), slower extension-decrease rates during the linear phase, and longer times to complete reversal after initiation of the final drop (t_1_) (**Fig. S6**). These results indicate that the mutation selectively impairs progression beyond lagging-strand unwinding into the shortening phase that results from leading strand unwinding. Consistent with this model, in the ensemble assay the R447A K448A mutant affected activity on the *lag gap* substrate to a greater extent than on the *lead overhang* substrate (**Fig. 1F**). Moreover, the mutant retained fork reversal activity on the substrate with both binding sites, which suggests that unwinding of both nascent strands is sufficient for fork reversal and that the mutated residues are involved in the leading strand destabilization step.

Together, these results support a reversal mechanism in which FBH1 first unwinds the lagging strand and, through its interaction with the fork junction, subsequently destabilizes the leading strand fork junction to promote unwinding of the nascent leading strand. This sequential unwinding exposes complementary parental strands, which reanneal passively due to their intrinsic complementarity. As a result, fork reversal by FBH1 is strongly dependent on conditions that permit spontaneous strand pairing. Increased opposing force stretches the fork and inhibits both nascent duplex melting and parental strand annealing, thereby reducing the efficiency of reversal at 4 pN. This behavior contrasts with SNF2-family remodelers, which presumably actively drive fork reversal and remain effective under high mechanical loads, indicating that FBH1 operates through a fundamentally distinct, force-sensitive mechanism.

### Products of SCF^FBH1^-mediated fork reversal are distinct from four-way junctions

The single-molecule data and structure of SCF^FBH1^ bound to the lagging strand template(*23*) imply that FBH1 does not generate a four-way junction during fork reversal. To test this, we visualized products of SCF^FBH1^-mediated remodeling of a stalled fork–like DNA substrate using atomic force microscopy (AFM). The substrate matches that used in magnetic-tweezers experiments but was adapted for AFM imaging and contains two dsDNA arms of unequal length joined by a short hairpin and an 8-nt single-stranded gap on the lagging strand at the fork (**Fig. 2A**). Canonical fork reversal of this three-way junction allows branch migration along two 495-bp migratable segments before reaching a 1,524-bp non-migratable region on the leading strand and a 20-bp non-migratable tip on the lagging strand (**Fig. 2B**). Direct measurement of the contour length of bare DNA molecules yielded a rise per base pair of 0.31 ± 0.01 nm bp^-1^ (mean ± SD), consistent with B-form DNA and in agreement with previous AFM measurements(*43, 44*) (**Fig. S7**). As a control for four-way junction detection, RecG was incubated with the substrate in the presence of ATP. This produced canonical T-shaped Holliday-junction structures (**Fig. 2C**) whose measured arm lengths corresponded to the expected 495-bp and 1,524-bp segments (**Fig. 2D**), confirming reliable identification of four-way intermediates in our assay.

**Fig. 2.**
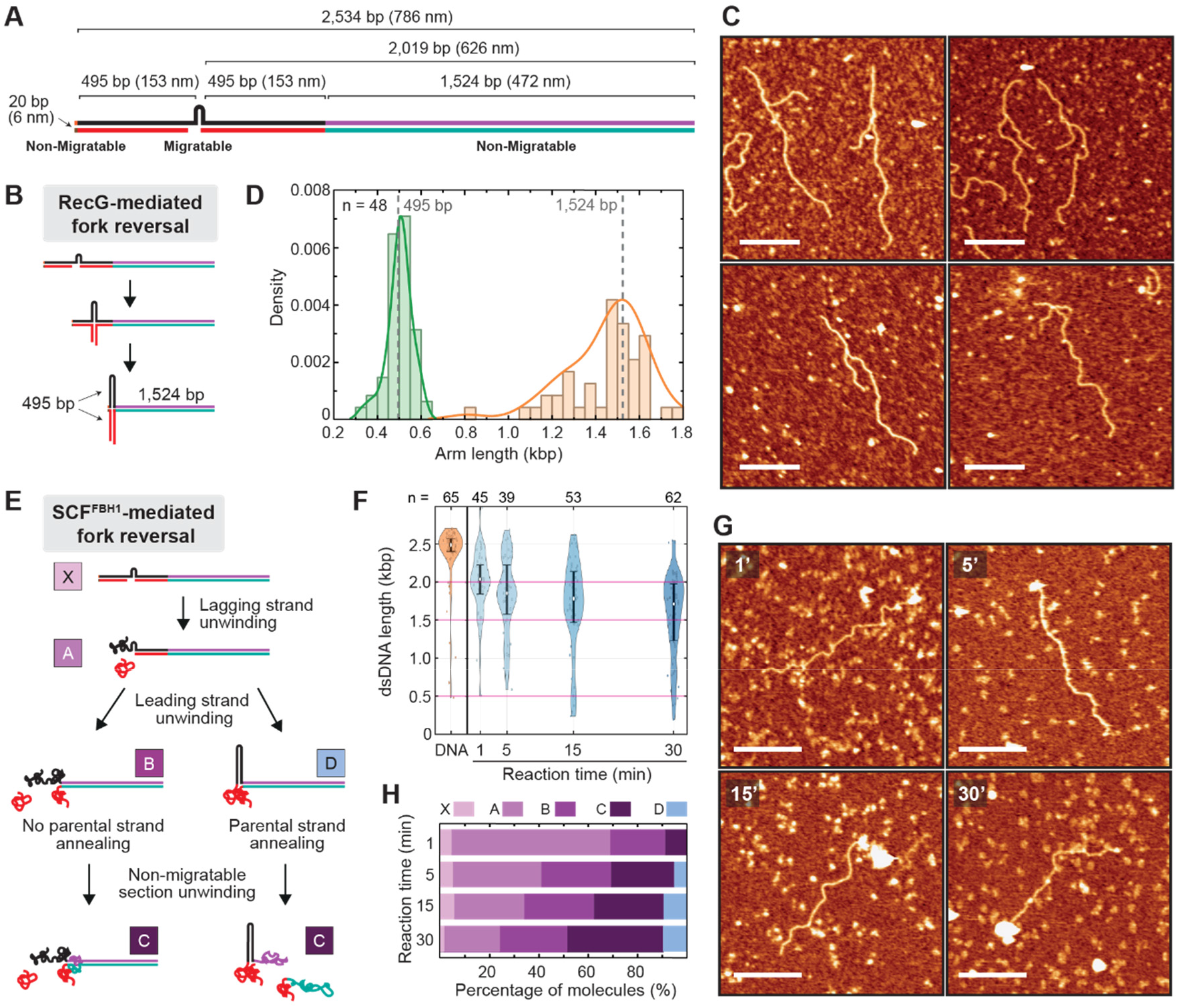
SCF^FBH1^-mediated fork reversal generates ssDNA-containing intermediates distinct from four-way junctions. **A**. Schematic of the AFM substrate. **B**. Schematic of expected DNA intermediates and products of RecG-mediated fork reversal. **C**. Representative AFM images of products generated by RecG on the fork substrate. Scale bar: 200 nm. **D**. Quantification of DNA arm lengths in RecG reaction products. Green and orange histograms correspond to the short and long arms, respectively. Solid curves show kernel density estimates. Dotted lines indicate the expected arm lengths, and *n* represents number of molecules observed. **E**. Proposed reaction pathway for SCF^FBH1^-mediated fork reversal. Lettered intermediates correspond to the classes quantified in panel H. **F**. Distribution of dsDNA segment lengths as a function of reaction time, with n specifying the number of molecules (observations). **G**. Representative AFM images of products formed by SCF^FBH1^ at the indicated time points on the fork substrate. Scale bar: 200 nm. **H**. Time-dependent distribution of SCF^FBH1^ reaction intermediates.

In ATP-free SCF^FBH1^ reactions, measurement of DNA arms emerging from the SCF^FBH1^-bound position yielded lengths of 152 ± 4 nm (494 ± 13 bp) and 647 ± 5 nm (2,103 ± 16 bp) (**Fig. S7**), consistent with specific binding at the fork junction. To test whether reversal proceeds through the mechanism inferred from magnetic-tweezers and ensemble fork-reversal experiments (**Fig. 2E**), SCF^FBH1^, DNA substrate, and ATP were incubated for 1–30 min before deposition. The predominant species consisted of a continuous dsDNA segment ending in a dense nucleoprotein cluster (**Fig. 2F-G**), consistent with protein accumulation on ssDNA generated by unwinding. The length of dsDNA extending from this cluster decreased progressively with reaction time **(Fig. 2F-G**). In the 1-min sample, a prominent population centered near ∼2 kbp corresponds to a product expected from lagging strand unwinding.

Molecules were classified into reaction intermediates according to dsDNA length and whether the nucleoprotein cluster occupied a terminal or internal position (**Fig. 2E-H, Fig. S8, Sup. Methods**). Lagging strand unwinding predominated at the earliest time point, whereas later samples showed increasing leading strand unwinding, extending beyond the migratable region into the non-migratable leading strand segment. Unwinding occurred either without or with parental strand annealing (classes C and D, respectively). The fraction of molecules showing annealed parental strands increased at later time points (**Fig. 2H**), consistent with progressive strand reannealing. The persistence of molecules that lack parental annealing despite extensive unwinding is consistent with continued FBH1 association with generated ssDNA. We did not observe accumulation of canonical four-way junction structures comparable to those formed by RecG at any time point, consistent with a mechanism involving sequential unwinding of lagging and leading strands that do not associate to form a stable Holliday-junction product.

### Fork reversal by SCF^FBH1^ does not involve nascent strand annealing

To further investigate the nature of the intermediates and products generated by SCF^FBH1^, we developed a fluorescence-based fork reversal assay to monitor the rates of annealing of parental and nascent strands separately (**Fig. 3, Fig S9**). In the assay, parental (**Fig. 3A**) or nascent (**Fig. 3B**) strands were labeled with a fluorescein (FAM)–black-hole quencher (BHQ1) pair, and fork reversal measured by fluorescence quenching as the strands anneal. If a Holliday junction is formed, the rates of parental and nascent strand annealing would be the same. In the presence of SCF^FBH1^, however, we observed that the rate of annealing of parental strands occurred faster and was more dependent on protein concentration than the nascent strands (**Fig. 3C, Fig. S10**). The rate of nascent strand annealing in the presence of SCF^FBH1^ was the same as their rate of spontaneous annealing (**Fig. 3D**), consistent with a model in which FBH1 unwinding of leading and lagging arms leads to a rapid zippering of parental duplex followed by slower, spontaneous annealing of the nascent strands. In contrast, HLTF annealed parental and nascent strands at the same rate (**Fig. 3D**), as expected with Holliday junction formation, providing further evidence that the mechanism of reversal is different between FBH1 and the SNF2-family remodelers.

**Fig. 3.**
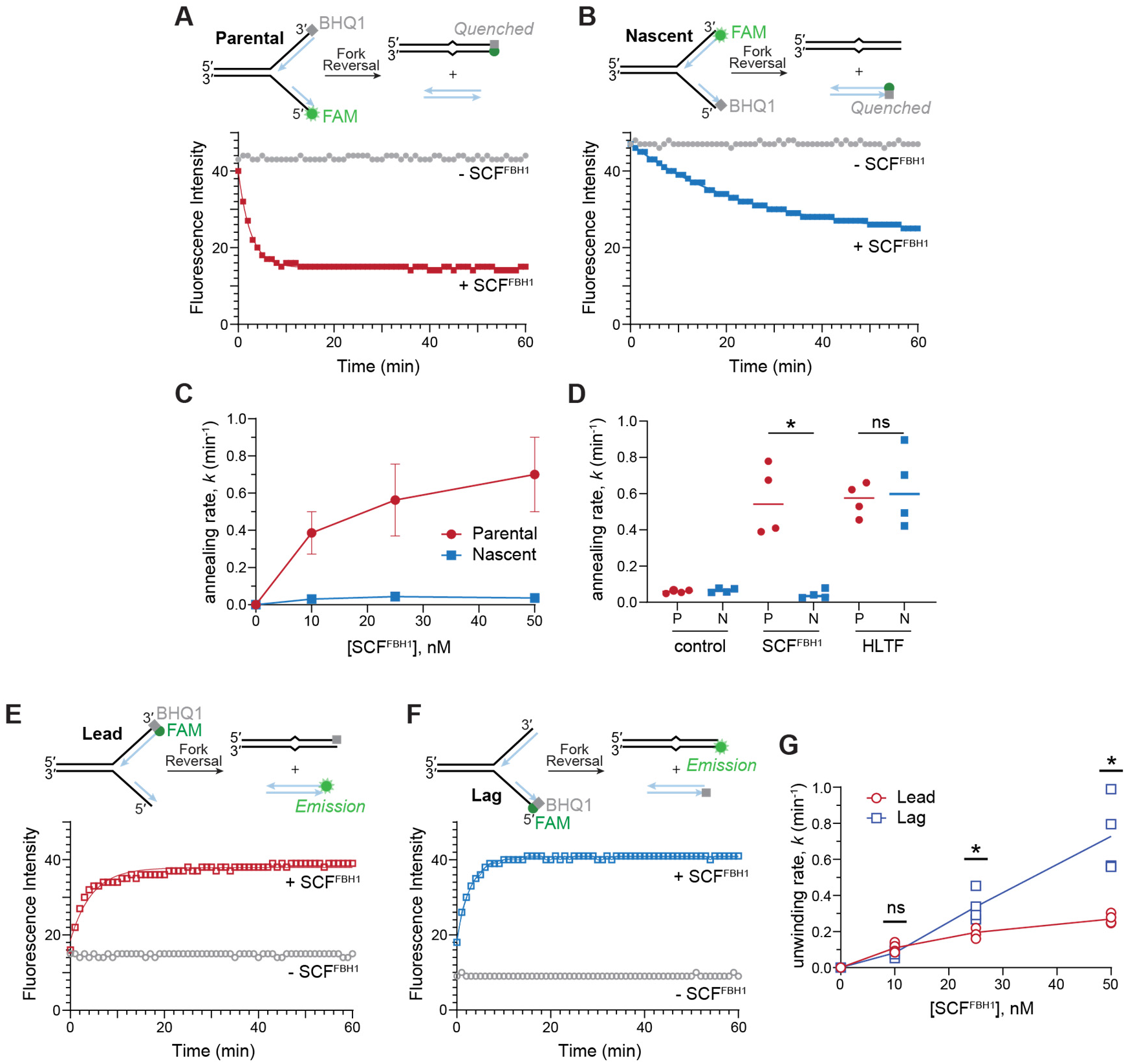
Fork reversal by SCF^FBH1^ does not involve nascent strand annealing. **A**,**B**. Representative traces from annealing of parental (A) or nascent (B) strands that contain a FAM-BHQ1 pair. Schematics of the fluorescence-based fork reversal assays are shown above the fluorescence traces. **C**. Rates of parental and nascent strand annealing during fork reversal at various SCF^FBH1^ concentrations (mean ± SD, n=4). **D**. Annealing rates of parental (P) and nascent (N) strands either free in solution prior to substrate formation (control) or as part of a fork substrate in the presence of SCF^FBH1^ or HLTF. Lines represent mean values, and significance was determined using an unpaired t-test with Welch’s correction (*, p=0.012; ns, not significant). Data for SCF^FBH1^ is the same as that shown in panel C (25 nM). **E**,**F**. Representative fluorescence traces of unwinding leading (E) or lagging (F) strands. **G**. Quantification of leading and lagging arm unwinding rates during fork reversal at various SCF^FBH1^ concentrations (mean ± SD, n=4). Significance for each concentration was determined using an unpaired t-test with Welch’s correction (*, p=0.0347 (25 nM) and p=0.0203 (50 nM); ns, not significant).

Because FBH1-mediated fork reversal is stimulated by nascent strand unwinding, we used the fluorescence assay to examine unwinding of each nascent strand. Fluorescence was monitored as a function of time from forks containing a FAM–BHQ1 pair on either leading (**Fig. 3E**) or lagging (**Fig. 3F**) arms. At higher concentrations of SCF^FBH1^, we observed faster rates of lagging strand unwinding compared to those of the leading strand (**Fig. 3G**), consistent with lagging strand translocation facilitating destabilization of the leading duplex(*23*). Interestingly, unwinding rates reached parental annealing rates only at the highest protein concentration tested (**Fig. 3C,G**). Since the fluorophore is present at the end of the duplexes to be unwound, these data suggest that the parental arms quickly anneal as the nascent strands are removed from the vicinity of the junction.

### Two SCF^FBH1^ complexes can bind to a single fork

Unwinding both leading and lagging strands raises the question of stoichiometry of SCF^FBH1^ at the fork. We used electrophoretic mobility shift assays (EMSAs) to examine SCF^FBH1^ binding to immobile fork structures containing ssDNA binding regions on the lagging template (*lag gap*), the nascent leading strand (*overhang*), or both (*gap+overhang*) (**Fig. 4A**). As previously observed(*23*), multiple protein-DNA bands were evident in each substrate, consistent with two SCF^FBH1^ complexes bound to fork (**Fig. 4A**). Although the highest affinity was observed for the forks containing a lag gap, two protein-DNA bands were observed regardless of the number of ssDNA binding sites, suggesting that at higher concentrations the protein displaces the nascent strands. We also observed two SCF^FBH1^ complexes per fork in mass photometry and negative stain EM experiments with the *gap+overhang* fork (**Fig. 4B-C**). Mass photometry showed four peaks corresponding to (i) buffer, (ii) FBH1, (iii) SCF^FBH1^ bound to DNA, and (iv) two SCF^FBH1^ complexes bound to DNA, the latter of which accounted for 10.6 ± 4.8% of the total (**Fig. 4B, Fig. S11**). Negative stain EM, which yielded ∼15,000 distinct SCF^FBH1^ particles that we sorted into nine 2D classes, showed five classes containing two colocalized SCF^FBH1^ molecules (**Fig. 4C**). Because the relative orientation of the two SCF^FBH1^ particles was not consistent among these classes, it is unlikely that the protein is forming a dimeric interaction. Rather, these data, together with the low abundance of the 2×SCF^FBH1^•DNA complexes from mass photometry, suggest that two SCF^FBH1^ complexes are bound independently to a single fork and that simultaneous binding is not essential for fork reversal.

**Fig. 4.**
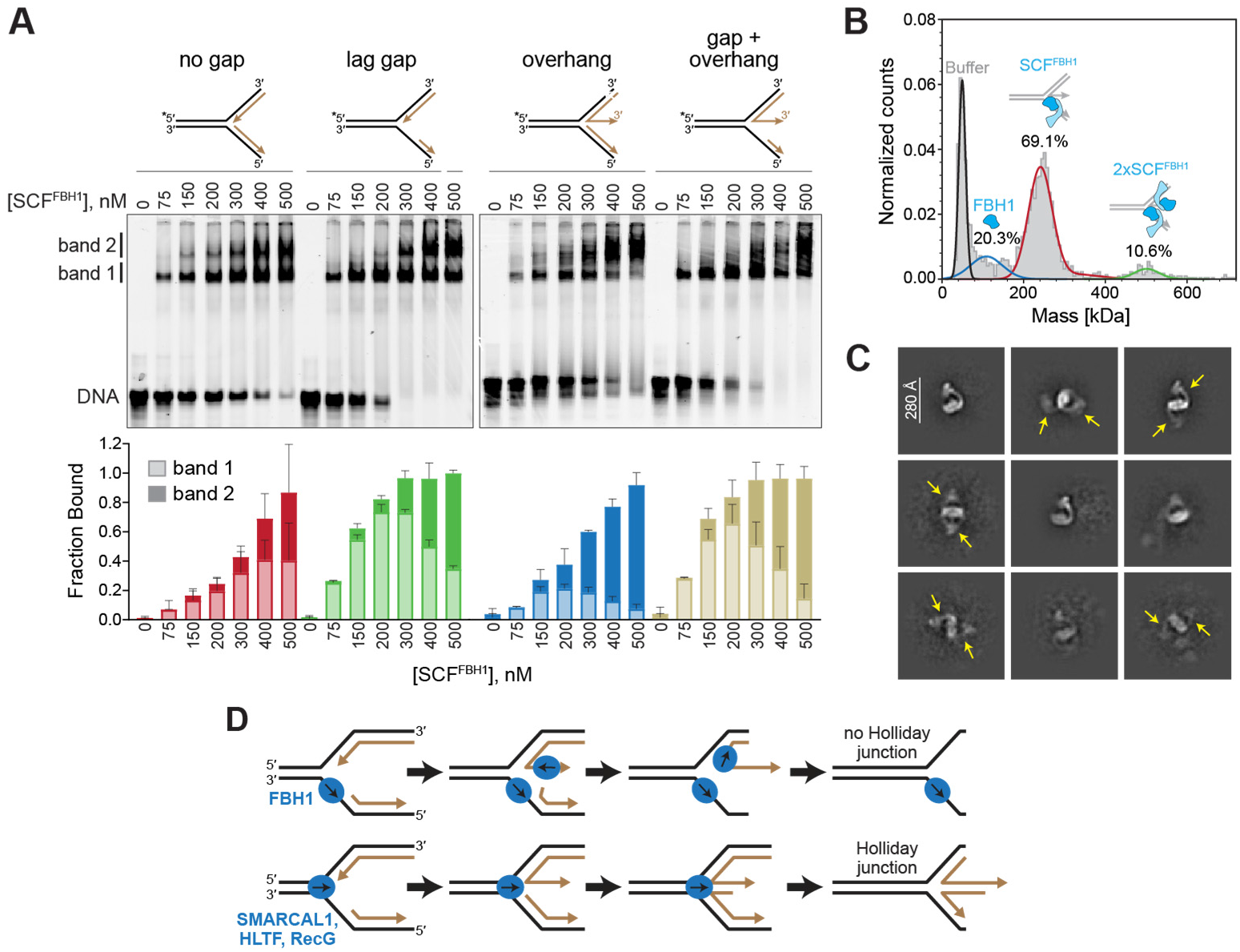
Two SCF^FBH1^ complexes can bind to a single fork. **A**. Representative native PAGE EMSA of SCF^FBH1^ binding to model fork substrates. Quantification of bands 1 and 2 (defined to the left of the gel) from multiple experiments is shown below (mean ± SD, n=3). **B**. Representative mass distribution of SCF^FBH1^ bound to the *gap+overhang* substrate with Gaussian curves fitted for each peak. Histogram peaks correspond to the number of binding events (counts) at a certain mass, and Gaussian curves reflect the average mass for each peak. Percentages reported are averages of the proportion of counts for each peak compared to the total number of raw counts, excluding counts attributed to buffer (n=3). **C**. Negative stain EM 2D class averages of SCF^FBH1^ bound to the same *gap+overhang* substrate as in panel B. Arrows indicate the position of multiple SCF^FBH1^ complexes in a single class. **D**. Proposed model of fork reversal by FBH1 (top) or SNF2-family remodelers (bottom).

## Discussion

The results presented here identify a mechanistically distinct pathway of replication fork reversal driven by helicase-mediated strand separation instead of duplex translocation (**Fig. 4E**). Rather than coupling fork reversal to dsDNA translocation and active annealing of parental and nascent strands, SCF^FBH1^ acts through a sequential unwinding reaction. In this pathway, the helicase first associates with and translocates along the lagging strand template, which displaces any nascent lagging strand in its path and promotes destabilization of the leading strand duplex, ultimately leading to unwinding of the nascent leading strand. This second step of leading strand destabilization is highly sensitive to opposing force and requires FBH1 interaction with the fork junction, indicating that it represents a discrete mechanistic transition rather than a simple continuation of lagging strand unwinding.

The strong force dependence observed in our single-molecule experiments provides insight into the physical basis of this transition. Lagging strand unwinding persists, and even increases in frequency, under higher tension, whereas progression to full reversal is strongly inhibited. This asymmetry indicates that the principal energetic barrier in the reaction is not helicase translocation itself, but destabilization of the junction resulting in exposure of ssDNA on the nascent leading strand. This interpretation is further supported by ensemble fork reversal assays showing that a leading strand 3′-ssDNA overhang strongly stimulates reversal. In our magnetic-tweezers substrates, the lagging strand contains a ssDNA gap that permits direct helicase loading, whereas the nascent leading strand remains base-paired up to the fork junction. SCF^FBH1^ therefore must first destabilize this duplex before productive engagement of the leading strand can occur.

Mutational and structural observations suggest that this transition involves direct protein–DNA interactions at the fork junction. Residues R447 and K448 form part of a positively charged surface that contacts the leading strand branchpoint at the fork junction, and mutation of these residues selectively impairs fork reversal without affecting helicase activity(*23*). These interactions, combined with translocation along the lagging strand, likely generate local strain at the fork junction that promotes duplex destabilization and exposes ssDNA on the nascent leading strand. Once a sufficient amount of ssDNA is exposed, loading and translocation of SCF^FBH1^ on the nascent leading strand can proceed, allowing parental strands to reanneal through intrinsic complementarity. In this view, SCF^FBH1^ does not actively rewind parental duplex DNA, but instead creates the conditions for reversal by sequentially removing the nascent strands. Consistent with this model, negative-stain EM and mass photometry analyses indicate that two SCF^FBH1^ complexes can bind simultaneously to a single fork substrate, raising the possibility that two helicase molecules could act on opposite arms of the fork during sequential unwinding. One molecule engages the lagging strand at the junction while another binds the 3′-end of the nascent leading strand once it becomes exposed.

Our data indicate that this mechanism produces an intermediate that differs fundamentally from the canonical reversed fork generated by SNF2-family remodelers SMARCAL1, HLTF, ZRANB3, and RecG. These enzymes act as dsDNA translocases that couple parental strand annealing with nascent strand pairing to generate a symmetric four-way junction. In contrast, SCF^FBH1^ generates an asymmetric, ssDNA-rich intermediate distinct from a Holliday junction. Fork reversal in this system therefore appears to arise from nascent strand separation followed by passive parental strand reannealing, rather than from an active strand annealing motor. This interpretation is consistent with previous single-molecule FRET experiments showing that FBH1 undergoes repetitive shuttling on ssDNA gaps adjacent to fork junctions before extensive unwinding occurs(*37*). Although those substrates lacked homologous parental arms and therefore could not undergo reversal, the observed shuttling is consistent with the helicase repeatedly engaging the junction while translocating on the gap. Based on the cryo-EM structure of SCF^FBH1^ bound to a fork(*23*), junction-bound translocation would destabilize the adjacent leading strand duplex.

Genetic studies also indicate that FBH1-dependent fork reversal follows a pathway distinct from those driven by SNF2 remodelers. RAD54L has been implicated in two reversal pathways with different mechanistic requirements. In the HLTF/SMARCAL1 pathway, RAD54L is required but its branch migration activity is dispensable, whereas RAD54L branch-migration activity is essential for FBH1-dependent reversal (*39*). One explanation for this difference is that the intermediate produced by FBH1 is structurally different from the canonical reversed fork generated by SNF2 remodelers. Our results provide a mechanistic basis for this idea by showing that SCF^FBH1^ produces an ssDNA-rich intermediate that may require RAD54L-dependent branch migration to extend or stabilize a regressed arm.

Additional support for structural differences between reversal intermediates comes from studies of fork protection. Reversed forks generated by SNF2 remodelers are stabilized by protection factors including BRCA1, BRCA2, FANCD2, and BOD1L, which limit nucleolytic degradation of the regressed arm(*38*). In contrast, FBH1-dependent reversal relies on a distinct set of protection factors(*38*). A plausible explanation is that the ssDNA-rich intermediate produced by SCF^FBH1^ presents different structural vulnerabilities and therefore recruits or requires a separate protective network.

Recent work on the nuclease–helicase DNA2 further supports the idea that distinct fork-reversal intermediates have different biological consequences. DNA2 restricts recombination-based replication restart from persistent reversed forks, and DNA2 depletion leads to extensive RPA-coated ssDNA and checkpoint activation(*45*). Notably, this RPA–ssDNA accumulation is strongly reduced by depletion of FBH1 but not by depletion of SNF2-family remodelers. Consistent with this, FBH1 inactivation has previously been reported to reduce ssDNA accumulation during replication stress, and mutation of the FBH1 helicase domain similarly diminishes ssDNA formation at stalled forks(*22*). Together, these observations support a model in which FBH1-mediated fork remodeling generates intermediates with elevated ssDNA content that may be particularly prone to recombination-dependent restart or pathological processing under persistent stress.

Emerging evidence suggests that helicase-driven fork remodeling may represent a broader mechanistic class. For example, the Ski2-like helicase ASCC3 has been shown to unwind DNA at stalled replication forks, promoting fork reversal and resulting in the accumulation of RPA-coated ssDNA during replication stress(*46*). Although the molecular pathway underlying ASCC3-dependent remodeling remains unresolved, these findings support the idea that fork reversal should not be viewed as a single mechanistic process. Instead, distinct classes of DNA motors may remodel stalled forks through fundamentally different physical mechanisms, with diverse structural outcomes.

Several questions about the distinct FBH1 fork reversal pathway remain. In particular, the extent to which parental strand reannealing proceeds passively after nascent-strand unwinding would depend on the presence and potential interaction with other proteins at stalled forks. We also do not know the precise architecture of the FBH1-mediated reversed fork and whether it facilitates a recombination-independent restart pathway. Moreover, the fact that FBH1 generates ssDNA but not a four-way junction raises the possibility that the nascent strand degradation assay may not be an accurate proxy for fork reversal in all cases. Defining the precise regressed DNA intermediates and products will be important for understanding how helicase-driven fork remodeling is integrated into cellular responses to replication stress.

## Materials and Methods

### Protein expression and purification

SCF^FBH1^ was expressed and purified as described(*23*) with minor modifications. Briefly, proteins were produced in baculovirus-infected Sf9 insect cells from a 438-C expression vector containing human 6xHis-MBP-FBH1 isoform 4, SKP1, CUL1, and RBX1. The supernatant from lysed cells was subjected to Ni-NTA (HisPur, Thermo Fisher) affinity resin equilibrated in Buffer N (20 mM HEPES pH 7.5, 500 mM NaCl, 5% glycerol, 10 mM imidazole, 0.5 mM TCEP), and SCF^FBH1^ proteins eluted with 200 mM imidazole in Buffer N. Ni-NTA elutions were dialyzed overnight at 4°C against 0.1 mg/ml TEV protease in Buffer H (20 mM HEPES pH 7.5, 100 mM NaCl, 5% glycerol, 0.5 mM TCEP), followed by centrifugation at 4,000 ×g for 10 min. The supernatant was passed through Ni-NTA resin, washed with 20 mM imidazole/Buffer H, and proteins further purified on a Heparin HiTrap column (Cytiva). Fractions containing SCF^FBH1^ were pooled, concentrated to 2.5 mg/mL, and flash frozen in liquid nitrogen. Human HLTF and *Thermatoga maritima* RecG was expressed and purified as previously described(*47, 48*).

### DNA substrate preparation

#### Magnetic tweezers

The MT fork substrate used in the reversal/branch migration assay was adapted from our previously published design(*23*) with minor modifications. The substrate consists of two identical dsDNA fragments arranged in an inverted orientation and connected by a short hairpin, forming a three-way junction capable of branch migration. To prevent spontaneous branch migration, a single base-pair mismatch was introduced at the base of the hairpin. In this configuration, the substrate contains an 8-nt ssDNA gap on the nascent lagging strand, which is required for SCF^FBH1^ fork reversal activity. The structure is ligated to a 997-bp digoxigenin-labeled fragment and a 152-bp biotin-labeled fragment that serve as immobilization handles. For the complementary unwinding assay, which lacks parental-strand annealing, a gapped DNA substrate was used. This substrate consists of a 6,610-bp dsDNA molecule containing a 63-nt ssDNA gap that allows protein loading, positioned 445 bp from the surface. The construct is ligated to two highly labeled DNA fragments used as handles—a 1,003-bp fragment labeled with digoxigenins and a 144-bp fragment labeled with biotins. Detailed procedures for substrate construction are provided in the Supplementary Methods.

#### Atomic force microscopy

The AFM fork substrate used in the reversal/branch migration was based on the magnetic tweezers construct, with modifications suitable for AFM imaging. The substrate consists of two dsDNA arms of unequal length joined by the short hairpin, forming a three-way junction. This configuration supports branch migration along two 495-bp migratable segments terminating at non-migratable regions of 1,524 bp on the leading strand and 20 bp on the lagging strand. An 8-nt single-stranded gap is positioned on the lagging strand adjacent to the fork to facilitate protein loading, and a single base-pair mismatch is introduced at the base of the hairpin to prevent spontaneous branch migration. Detailed procedures for substrate construction are provided in the Supplementary Methods.

#### Ensemble biochemistry

Oligodeoxynucleotides used in this study are listed in **Table S1**, and annealed strand combinations for each substrate specified in **Table S2**. Oligos were initially annealed from 95 °C to 25 °C over 70 min in 1× SSC buffer (15 mM sodium citrate pH 7.0 and 150 mM NaCl). All fork substrates were formed by first annealing the leading and lagging strand arms separately, followed by annealing the duplexed arms at 37°C for 30 min. Fluorescence fork reversal assay substrates with 5′-6-carboxyfluorescein (FAM)- and 3′-Black Hole Quencher 1 (BHQ1)-labeled oligonucleotides were generated with a 1:1.1 molar ratio of labeled:unlabeled or FAM-labeled:BHQ1-labeled for each duplex. For gel-based fork reversal assay substrates, *oligo48* was first 5′-labeled by incubating with ^32^P-γATP and polynucleotide kinase (NEB) at 37°C for 60 min, followed by heat inactivation at 65 °C for 10 min and purification via a G-25 column (GE Healthcare). The ^32^P-labeled leading (*48/50* or *48/50_10polyT*) and lagging (*52/53* or *52/53_5gap*) halves of the fork were annealed separately, followed by incubation of unlabeled and labeled duplexes in a 2:1 molar ratio. ^32^P-labeled substrates were gel purified using a 6% 0.5× TBE DNA Retardation Gel (Invitrogen), from which bands were excised and electroeluted in 0.5× TBE. Substrates were concentrated using 10K Amicon-Ultra 0.5 mL centrifugal filters and stored at -20°C. FAM-labeled fork substrates used for binding studies were annealed using a 1.2-fold molar excess of unlabeled duplex. The substrates for mass photometry and EM were annealed at a 1:1 molar ratio.

### Magnetic tweezers

Experiments were performed using a custom-built instrument similar to previously described setups(*49*–*51*). Briefly, a pair of vertically aligned permanent NdFeB magnets was positioned above a flow chamber mounted on an inverted microscope. DNA molecules were tethered between the chamber surface and 1-µm-diameter MyOne C1 superparamagnetic beads (Invitrogen). Vertical translation of the magnets controlled the stretching force applied to the DNA. Bead positions were recorded at 120 Hz using a CCD camera and a high-magnification oil-immersion objective. DNA extension was determined by video-based particle tracking and reconstruction of bead height from focal plane calibration. Forces were calculated from the variance of lateral Brownian fluctuations of individual beads using the equipartition method(*49*).

Flow chambers were assembled by sandwiching a single parafilm spacer between two glass coverslips to define the flow channel; the upper coverslip contained inlet and outlet holes for buffer exchange. The magnets (Supermagnete) were separated by a 1-mm gap, enabling forces up to approximately 7 pN on 1-µm beads. Flow chambers with polystyrene-coated lower coverslips were functionalized by passive adsorption of anti-digoxigenin antibodies (100 ng µl^−1^, overnight at 4 °C) and subsequently passivated with 1 mg ml^−1^ bovine serum albumin in PBS. DNA–bead complexes were prepared by incubating biotinylated DNA with 1 µm streptavidin-coated magnetic beads and introduced into the chamber, allowing the digoxigenin-labelled DNA end to bind to the surface-immobilized antibodies. Unbound beads were removed by buffer flushing. The procedure followed previously described methods(*50, 52*).

Fork reversal assays were performed using a stalled replication fork-like DNA substrate, whose DNA extension was monitored continuously for 90 min at room temperature (21ºC). Prior to initiating the reactions with protein and ATP, flow chambers containing immobilized DNA molecules were equilibrated with the corresponding reaction buffer for each protein. Reactions were initiated by introducing 40 nM SCF^FBH1^ in FBH1 buffer (20 mM Tris–HCl (pH 8.0), 50 mM NaCl, 2 mM ATP, 5 mM MgCl_2_, and 1 mM DTT). HLTF reactions were initiated by introducing 40 nM HLTF in HLTF buffer (40 mM Tris–HCl (pH 7.5), 50 mM NaCl, 1 mM ATP, 2 mM MgCl_2_, and 2 mM DTT). Unwinding assays shown in **Fig. S3** were performed using the gapped DNA substrate. Flow chambers containing immobilized DNA molecules were equilibrated with FBH1 buffer. Reactions were initiated by introducing 40 nM SCF^FBH1^ in FBH1 buffer.

### Atomic force microscopy

AFM imaging was performed in tapping mode in air at room temperature and low humidity using a Nanotec Electrónica S.L. instrument equipped with an PPP-NCH-W PointProbe-PLUS cantilever (204-497 kHz resonance frequency). Images were acquired with a scan size of 2 µm × 2 µm at a scan rate of 1 line s^−1^ and 500 points per line. Images were processed using WSxM freeware(*53*). For quantification of products, images from multiple mica areas were collected. DNA contour length under the imaging conditions was calibrated by imaging naked SRFL-AFM DNA in FBH1 buffer lacking ATP and depositing it directly onto mica. Contours of individual DNA molecules were manually traced using WSxM, and a Gaussian fit to the resulting length distribution was used to determine the reference contour length. The lengths of dsDNA regions within individual molecules were determined by tracing their contours using the same procedure. In the case of FBH1 the reaction intermediates were classified following criteria in **Fig. S8**.

SCF^FBH1^ reactions were performed in bulk at 37 °C using 12 nM SCF^FBH1^ and 0.7 nM DNA substrate in FBH1 buffer supplemented with MgCl_2_ to a final concentration of 10 mM. After the desired incubation time, the reaction mixture was deposited onto freshly cleaved mica and allowed to adsorb for 5 min at room temperature. The mica was then rinsed with Milli-Q water, dried under nitrogen, and imaged in air. For imaging of SCF^FBH1^ binding to DNA, the substrate was incubated with the protein for 5 min at room temperature in FBH1 buffer without ATP [5 nM SCF^FBH1^, and 0.5 nM DNA substrate] and directly deposited onto freshly cleaved mica. After 5 min adsorption, the mica was rinsed with Milli-Q water and dried under nitrogen prior to imaging. RecG reactions were performed in bulk at 37 °C using 20 nM RecG and 0.7 nM DNA substrate in RecG reaction buffer (25 mM TrisOAc pH 7.5, 150 mM KOAc, 5 mM Mg(OAc)_2_, 1 mM DTT, and 1 mM ATP). For these samples, freshly cleaved mica was pretreated with 0.25 mM NiCl_2_ for 5 min, rinsed with Milli-Q water, and dried prior to deposition of the reaction mixture. After 5 min incubation on the mica at room temperature, the surface was again rinsed with Milli-Q water, dried under nitrogen, and imaged in air.

### Ensemble fork reversal assays

Fluorescence-based fork reversal reactions contained varying concentrations (0–50 nM) of SCF^FBH1^ or HLTF, 10 nM labeled fork substrates (**Table S1 and S2**), 100 μg/mL BSA, and 2 mM ATP in either 40 mM Tris pH 8.0, 50 mM NaCl, 5 mM MgCl_2_, and 1 mM TCEP (SCF^FBH1^) or 20 mM Tris pH 7.76, 10 mM KCl, 2 mM MgCl_2_, and 1 mM DTT (HLTF). Reactions were performed in a 384-well plate at 25°C or 37°C for 1 hour. Fluorescence intensity was measured with a Synergy H1 Hybrid Multi-Mode Microplate Reader (BioTek) using a fluorescein isothiocyanate filter set and quantified with Gen5 (BioTek) software. Fluorescence traces were analyzed using Prism10 (GraphPad) and fitted to a single exponential curve to determine rate constants.

Gel-based fork reversal reactions were performed at 21°C and contained 5 nM SCF^FBH1^ (WT or R447A K448A), 1 nM ^32^P-labeled fork substrate (**Table S1 and S2**), 2 mM ATP, 100 μg/mL BSA, and reaction buffer (40 mM Tris pH 8.0, 50 mM NaCl, 5 mM MgCl_2_, and 1 mM TCEP). At each time point, 10 μL of the reaction was stopped by the addition of Proteinase K (Sigma). Reactions were electrophoresed on an 8% 19:1 (acrylamide:bis-acrylamide) 1× TBE gel at 10 W for 1.5 hours and phosphorimaged on a Typhoon RGB imager (Cytiva). Band intensities were quantified with ImageQuantTL (Cytiva) and analyzed using Prism10 (GraphPad).

### DNA binding assays

Electrophoretic mobility shift assay reactions were performed on ice for 30 min and contained 0-500 nM SCF^FBH1^, 100 nM 5′-FAM labeled fork substrate (**Table S1 and S2**), and binding buffer (20 mM HEPES pH 7.5, 100 mM NaCl, 0.5 mM TCEP, and 0.02% NP-40). Samples were electrophoresed on a 5% 79:1 (acrylamide:bis-acrylamide) 1× TBE gel containing 5% glycerol (v/v) at 200 V for 1 hour. Gels were imaged with a Typhoon RGB imager (Cytiva), and band intensities quantified using ImageQuantTL (Cytiva) and analyzed using Prism10 (GraphPad).

For mass photometry measurements, 1 μM SCF^FBH1^ was incubated with fork substrate (**Table S1 and S2**) in a 1:1.2 protein:DNA molar ratio at 4°C for 30 min in Buffer B (20 mM HEPES pH 7.5, 150 mM KCl, 5 mM MgCl_2_, and 0.5 mM TCEP) prior to the addition of ATPγS (2 mM final concentration). Protein samples were diluted 40-fold in Buffer B before applying 5 μL of sample to 15 μL of Buffer B in each well on MassGlass^™^ UC slides, for a final protein concentration of 6.25 nM. Binding events were recorded using a Refeyn TwoMP Mass Photometer and quantified with Discover^MP^, using MassFerence^™^ P1 Calibrant to generate the calibration curve.

### Electron Microscopy

SCF^FBH1^ (44 nM) was incubated with the *gap+OH* fork substrate (**Table S1 and S2**) in a 1:1.2 protein:DNA molar ratio for 30 min on ice in Buffer B, followed by addition of ATPγS to a final concentration of 2 mM. Carbon film copper grids with 300 mesh (Electron Microscopy Sciences) were glow discharged for 2 min. 2.5 μL of sample was applied to the grid and incubated for 1 min. Excess sample was removed with filter paper and the grid stained with uranyl formate solution (0.7% w/v) with iterative cycles of applying and removing excess stain. Grids were screened and micrographs collected using a Thermo Fisher L120C TEM operating at 120 kV with a magnification of ×73,000 and dose rate of 647.34 e/nm^2^ using a Ceta-S detector. Particle picking and 2D classification were performed using cryoSPARC^45^.

### Statistical Analysis

Magnetic tweezers (MT) and atomic force microscopy (AFM) data processing and statistical analyses were performed using OriginPro. Time–extension traces from MT experiments were visually classified into the three predefined event types. For comparisons of kinetic parameters between the R446A-K447A mutant and wild-type (WT) proteins, data distributions were first assessed for normality. Depending on the outcome, either a two-sample t-test (for normally distributed data) or a Mann–Whitney U test (for non-normally distributed data) was applied. Parental strand closure rates in MT experiments were determined by linear regression of the final extension decrease observed in Type I events. Double-stranded DNA (dsDNA) contour lengths from AFM images were obtained by manual tracing using WSxM (*53*).

Statistical analyses in ensemble biochemistry experiments were performed using Prism10 (GraphPad). An unpaired, two-tailed Student’s t-test with Welch’s correction was applied to compare rates between two substrates. The number of replicates and individual *P*-values for each experiment are reported in the figure legends with a *P*-value of <0.05 reported as significant.

## Author contributions

Conceptualization: JMG, EMP, BFE, FMH

Methodology: JMG, EMP, CAR Investigation: JMG, EMP

Supervision: BFE, FMH

Writing—original draft: JMG, EMP, BFE, FMH

Writing—review & editing: JMG, EMP, CAR, BFE, FMH

## Data and materials availability

All data needed to evaluate the conclusions are present in the paper or the Supplementary Materials.

