## Supplementary Materials for "FBH1 Reverses Stalled Replication Forks via Sequential Unwinding of Nascent Strands"

**This file includes:**

Supplementary Text
Figs. S1 to S11
Tables S1 to S3
References (1 to 6)

### Supplementary Text

#### *Substrates for Single Molecule experiments*

The sequences of the oligodeoxyribonucleotides for used in magnetic tweezers (MT) and AFM experiments are described in Table S3.

MT migratable fork substrate. The design of a DNA substrate that mimics a stalled replication fork was fabricated following the protocol described in ref. (*1*) with slight modifications. Two identical dsDNA fragments arranged in an inverted orientation and connected by a short hairpin of 30 bp with a 4-nt loop ($dT_4$) were used to form a three-way junction capable of branch migration. A mismatch of 1 bp was included at the beginning of the hairpin to avoid spontaneous branch migration. In this case, the substrate contains an 8-nt ssDNA gap on the nascent lagging strand, strictly required for $SCF^{FBH1}$-driven reversal. The two 2-kbp dsDNA fragments were obtained by PCR amplification with Phusion High-Fidelity DNA Polymerase (Thermo Scientific) using the plasmid pSP73-JY0 (*2*) as template and oligonucleotides that include KpnI and PspOMI restriction sites in one side of the PCR fragment and a BbvCI site in the other side. After purification (QIAGEN), the dsDNA fragment that would be connected to the digoxigenin-labeled handle was digested with Nt.BbvCI restriction enzyme (NEB) creating a 13-nt 5'-overhang on one end. The dsDNA fragment that would be connected to the biotinylated handle was digested with Nb.BbvCI variant (NEB), creating an 18-nt 3'-overhang on one end. This strategy to generate both 3' or 5'-overhangs using nicking enzymes has been previously described (*3*, *4*). The 5'-overhang of 13 nt was annealed to 25-fold excess of oligonucleotide corresponding to the A-DIG side adaptor and the 3'-overhang of 18 nt was annealed with the A-BIO side adaptor by heating 10 min at 72°C and slowly cooling down to 42°C at a -0.1°C 25 $s^{-1}$ rate in annealing buffer (10 mM Tris-HCl pH 7.5, and 1 mM $MgCl_2$) followed by overnight ligation with T4 DNA Ligase (NEB). The complete 2-kbp–dsDNA branches were gel extracted and purified (QIAGEN). Then, the dsDNA fragment that would be connected to the digoxigenin-labeled handle was digested with PspOMI (NEB) and the dsDNA fragment that would be connected to the biotinylated handle was digested with KpnI (NEB), generating the overhangs to later ligate with the dsDNA handles. After digestion, the two branches were purified and mixed in equimolar ratio to subsequently perform the annealing between them through the DIG side adaptor and the BIO side adaptor, by heating as described above in the same annealing buffer. The oligonucleotide *250.Loop hairpin* was self-annealed by heating at 95°C for 5 min and cooling down to 20°C at a -1°C $min^{-1}$ rate in hybridization buffer (10 mM Tris-HCl pH 8.0, 1 mM EDTA, 200 mM NaCl, and 5 mM $MgCl_2$) to create a short dsDNA hairpin with a cohesive end compatible with the cohesive end formed once the DIG side adaptor and the BIO side adaptor were annealed between them. The dsDNA handles labeled with digoxigenins (997 bp) or with biotins (152 bp) were prepared by PCR including 200 µM final concentration of each dNTP (dGTP, dCTP, dATP), 140 µM dTTP and 66 µM Dig-11-dUTP or Bio-16-dUTP (Roche) using the plasmid pSP73-JY0 as template (*2*) followed by digestion with the restriction enzyme PspOMI or KpnI, respectively. In a final step, the two annealed branches were ligated overnight with the labeled handles together with a 10-fold excess of the self-annealed short dsDNA hairpin. The sample was ready for use without further purification. DNAs were never exposed to intercalating dyes or UV radiation during their production and were stored at 4°C with 1 mM EDTA pH 8.0.

MT non-migratable gapped substrate. This substrate consists of a central fragment of 6,610 bp ligated to two highly labeled DNA fragments, one with digoxigenins and the other with biotins, of 1,003 bp and 144 bp, respectively. The substrate is based on the pNLrep plasmid (kindly gifted by Prof. Dr. Ralf Seidel) (*3*) that has a DNA sequence with five closely-spaced BbvCI restriction sites. Nicking of one of the two strands with Nt.BbvCI enzyme results in the formation of short 15–16 nucleotides long fragments after heat denaturation, leaving a 63-nt ssDNA gap that allows protein loading, located at 445 bp from the surface. To fabricate this substrate, the pNLrep plasmid was digested with BamHI and BsrGI enzymes (NEB) followed by gel extraction. The 6,610-bp product was then digested with Nt.BbvCI. As described in ref. (*5*), to avoid reannealing of Nt.BbvCI cleavage products, a 100× excess of short oligonucleotides complementary to the four released Nt.BbvCI-fragments were added after Nt.BbvCI digestion and before inactivation of the enzyme for 20 min at 80°C. Following enzyme inactivation, the sample was cooled down to 40°C at a 1°C $min^{-1}$ rate. The 6,610 bp product with a 63 nt-gap was then purified with a PCR purification kit from QIAGEN. Highly-labeled handles were PCR-generated from the plasmid pSP73-JY0 using appropriate oligonucleotides as described above, including Dig-11-dUTP or Bio-16-dUTP, followed by restriction with BamHI or BsrGI, respectively. Finally, the labeled fragments were overnight ligated to the central part. DNAs were never exposed to intercalating dyes or UV radiation during their production and were stored at 4°C with 1 mM EDTA pH 8.0.

AFM substrate. The AFM fork substrate used in the reversal/branch migration matches that used in magnetic-tweezers experiments but was adapted for AFM imaging, containing two dsDNA arms of unequal length joined by the short hairpin. The three-way junction allows branch migration along two 495-bp migratable segments before reaching a 1,524-bp non-migratable region in the leading strand and a 20-bp non-migratable region in the lagging strand. The 8-nt single-stranded gap was also positioned on the lagging strand adjacent to the fork, and a mismatch of 1 bp at the beginning of the hairpin was also introduced to avoid spontaneous branch migration. The two dsDNA fragments of 0.5 and 2 kbp were obtained by PCR amplification with Phusion High-Fidelity DNA Polymerase using the plasmid pSP73-JY0 (*2*) as template and oligonucleotides that include a BbvCI site in one side. After purification, the dsDNA fragment that would be connected to the digoxigenin-labeled handle was digested with Nt.BbvCI restriction enzyme creating a 13-nt 5'-overhang on one end. The dsDNA fragment that would be connected to the biotinylated handle was digested with Nb.BbvCI variant, creating an 18-nt 3'-overhang on one end. The 5'-overhang of 13 nt was annealed with 25-fold excess of oligonucleotide corresponding to the A-DIG side adaptor and the 3'-overhang of 18 nt was annealed with the A-BIO side adaptor as described above, followed by overnight ligation with T4 DNA Ligase. The complete 0.5- and 2 kbp–dsDNA branches were gel extracted and purified. Next the two branches were mixed in equimolar ratio to subsequently perform the annealing between them through the DIG side adaptor and the BIO side adaptor, by heating as described above in the same annealing buffer. The oligonucleotide *250.Loop hairpin* was self-annealed as described for the MT substrate. In a final step, the two annealed branches were ligated overnight with a 5-fold excess of the self-annealed short dsDNA hairpin. The ligated final substrate was gel extracted and purified. DNAs were never exposed to intercalating dyes or UV radiation during their production and were stored at 4°C with 1 mM EDTA pH 8.0.

***Classification of FBH1 reversal reaction products from AFM images***

Images were classified based on the presence of protein clusters, their position, and the contour length of the dsDNA regions emerging from the cluster. Contour lengths were measured by manually tracing the molecule using WSxM Freeware (6). The classifications were based on a reaction model derived from magnetic tweezers and fluorescence experiments, in which the reaction proceeds through initial unwinding of the lagging strand followed by unwinding of the leading strand and, in some cases, parental strand annealing (**Fig. S8B**).

Structural categories. Molecules displaying a protein cluster were divided into two configurations: those in which two arms emerged from the cluster (internal cluster) and those in which a single dsDNA region emerged from the cluster (terminal cluster).

Class definitions.

- **Class X (naked DNA).** Molecules without any protein cluster. dsDNA contour length ≥ 715 nm (~2,306 bp).
- **Class A (lagging strand unwinding).** This class includes two types of molecules. Those with a terminal cluster and with dsDNA contour length ≥ 591 nm (~1,906 bp) and those with an internal cluster in which the dsDNA contour length of the long arm emerging from it ≥ 550 nm (~1,774 bp). This class corresponds to molecules in which unwinding is restricted to the lagging strand.

  The difference in the threshold between both configurations is introduced to ensure a clear separation from class D, in which parental strand reannealing is inferred. Assignment to class D requires a clear shortening of the leading strand arm, such that leading strand unwinding has occurred and reannealing is plausible. By using a less stringent cutoff in class A, we minimize the misclassification of partially unwound molecules as reannealed products. This conservative threshold also considers that contour length measurements may be affected by partial occlusion of dsDNA by the protein cluster.
- **Class B (partial leading strand unwinding).** Molecules with a terminal cluster and 469 nm ≤ dsDNA contour length < 591 nm (~1,513–1,906 bp). These molecules correspond to cases in which, after lagging strand unwinding, partial unwinding of the migratable region of the leading strand has occurred without parental strand annealing.
- **Class C (extensive leading strand unwinding).** Molecules with terminal cluster and contour length < 469 nm (~1,513 bp). These molecules correspond to cases in which, after unwinding of the lagging strand, unwinding of the leading strand has reached the non-migratable part of the substrate. Note, however that these molecules could also be the product of extensive unwinding followed by parental strand annealing, which would produce molecules with a cluster and a dsDNA region with contour length < 151 nm (487 bp), corresponding to the annealed parental strands (**Fig. S8B, lower part**). Thus, different mechanistic paths can lead to structurally similar products indistinguishable from AFM images. However, all the molecules with cluster and dsDNA contour length < 469 nm were classified as class C to avoid overestimating parental strand reannealing.

- **Class D (leading strand unwinding with parental strand reannealing).** Molecules with internal cluster and with the dsDNA contour length of the long arm emerging from it < 550 nm (~1,774 bp). These molecules are consistent with lagging and leading strand unwinding, resulting in parental strand annealing.

General considerations. The thresholds used do not correspond strictly to the expected physical lengths derived from bp-to-nm conversion. This is mainly due to partial occlusion of dsDNA regions by the nucleoprotein cluster and variability in contour tracing. Therefore, thresholds were chosen conservatively to minimize overestimation of leading strand unwinding and parental strand reannealing.

Parental strand annealing was only assigned to internal cluster configurations. However, some terminal cluster configurations with very short dsDNA arms are also compatible with parental strand reannealing. These cases were not separated as it is impossible to distinguish them in AFM images. Because of the conservative classification criteria used, the proportion of molecules in which parental strand reannealing has occurred likely represents a lower bound.

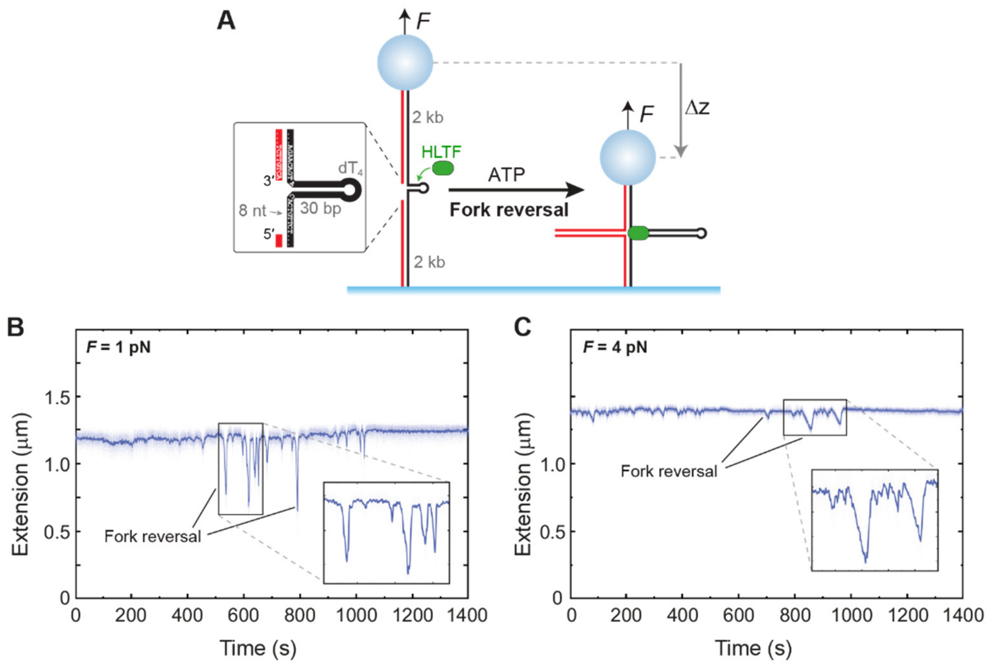

**Figure S1. HLTF reverses a fork substrate containing a ssDNA gap in the lagging strand. A**. Schematic of the magnetic tweezers assay. Fork reversal is detected as a measurable decrease in DNA extension. The inset shows details of the three-way junction design containing a ssDNA gap in the lagging strand. **B**,**C**. Time-extension traces showing DNA extension decreases and recoveries due to HLTF fork reversal activity against 1 pN (B) or 4 pN (C) opposing forces. Reactions contained 40 nM HLTF and 1 mM ATP.

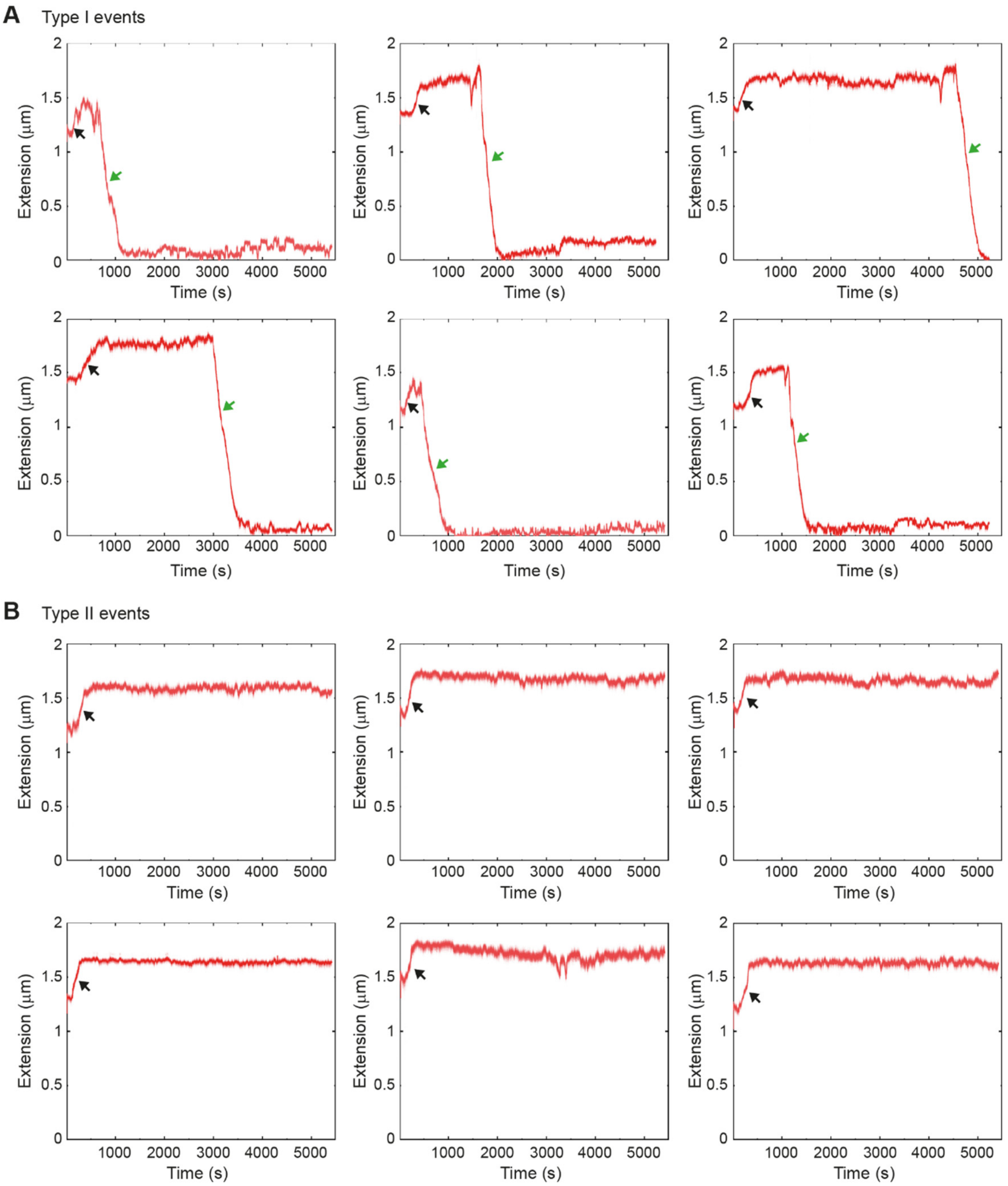

**Fig. S2. SCF$^{FBH1}$ reverses forks by unwinding both leading and lagging strands.** Representative time-extension traces of individual SCF$^{FBH1}$ activities on the fork substrate shown in Fig. 1A under 4 pN force. **A**. Type I events. DNA extension initially increases due to lagging strand unwinding (black arrows) and then decreases as leading strand unwinding and parental strand reannealing occur (green arrows). **B.** Type II events, which show only the initial extension increase from lagging strand unwinding (black arrows).

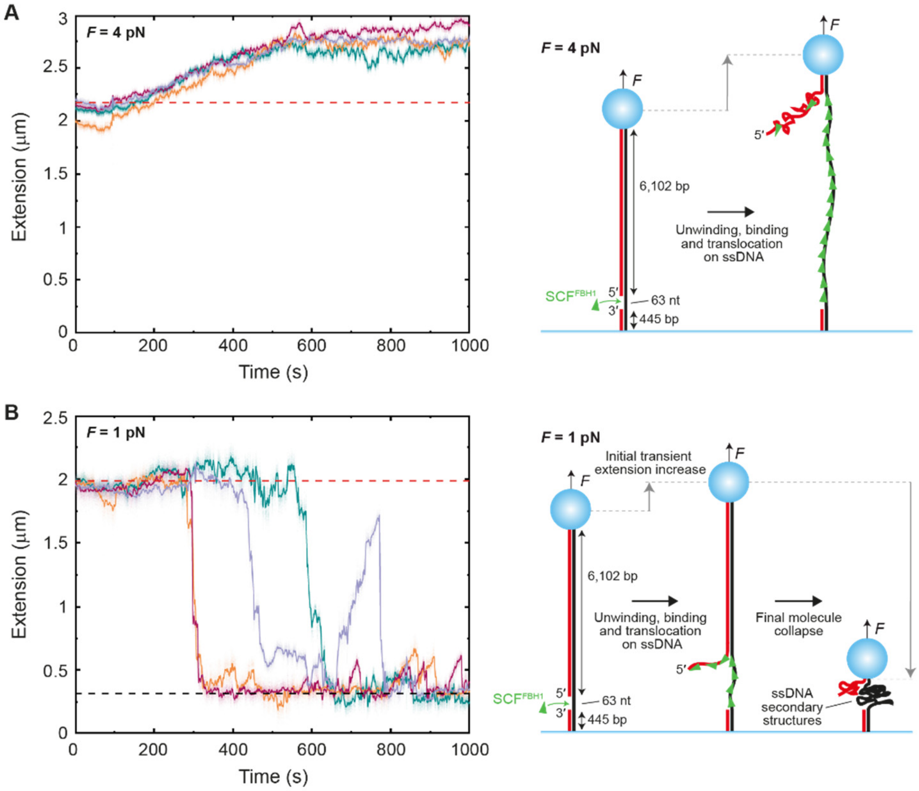

**Fig. S3. SCF$^{FBH1}$ unwinding activity in melting configuration. A**,**B**. Representative time-extension traces (*left*) and schematic illustrations (*right*) of SCF$^{FBH1}$ unwinding individual non-migratable DNA molecules containing a 63-nt ssDNA gap at forces of 4 pN (A) and 1 pN (B). In the experimental traces, the red dotted lines indicate the expected substrate extension prior to unwinding, and the black dotted line indicates the expected extension after unwinding. These reference lines are derived from force-extension curves for ssDNA and dsDNA. **A.** At 4 pN, unwinding leads to a net increase in extension, which can be attributed to protein binding and translocation on exposed ssDNA regions. **B**. At 1 pN, in a high proportion of events DNA extension transiently exceeds the initial level before ultimately decreasing. At this low force, unwinding can cause a transient increase in extension, followed by a collapse due to the formation of secondary structures in the resulting ssDNA.

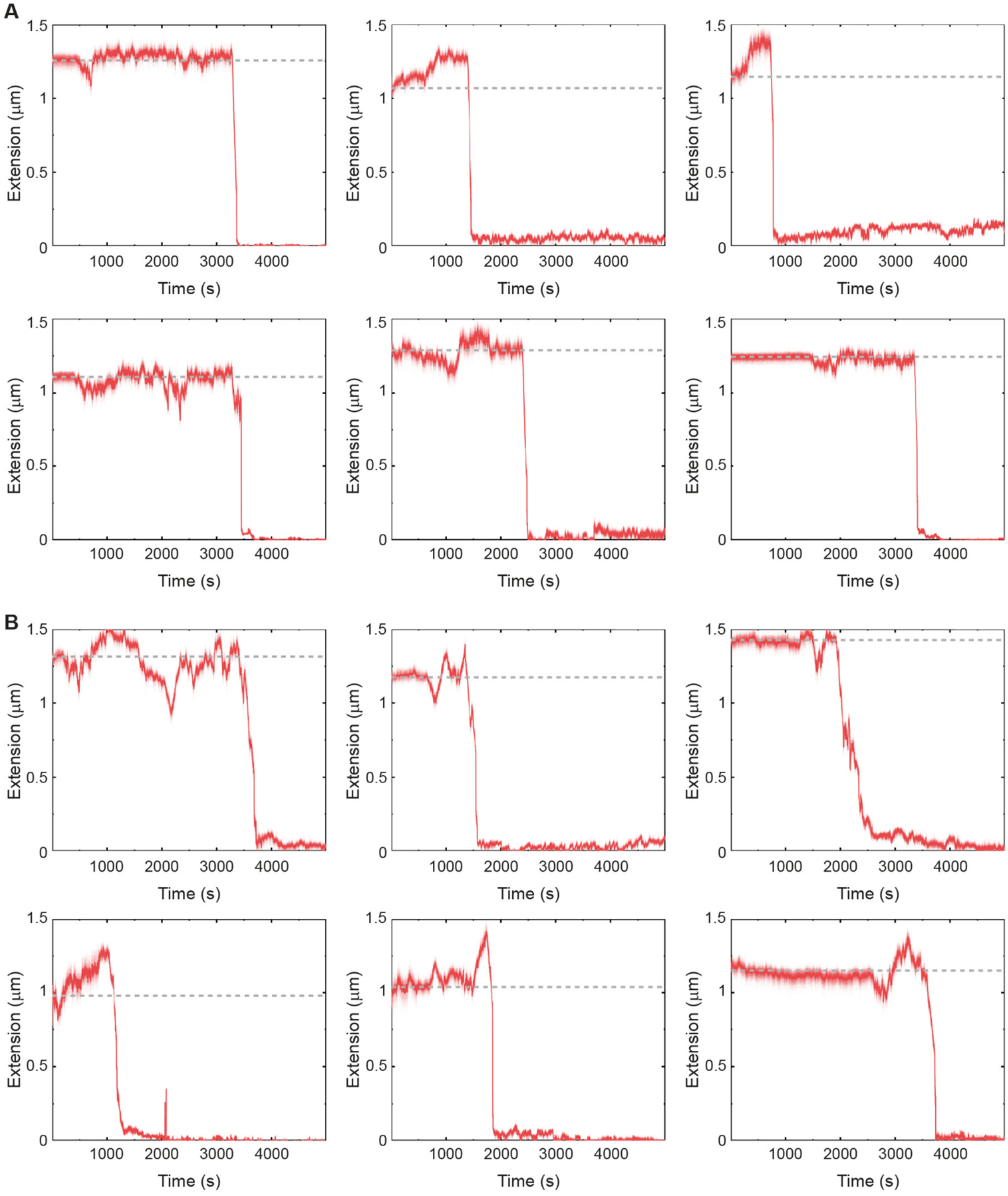

**Fig. S4. SCF$^{FBH1}$ activity on the fork substrate can induce transient extension increases above the initial level at 1 pN force.** Representative time-extension traces of individual wild-type (**A**) and R447A K448A mutant (**B**) SCF$^{FBH1}$ activities on the fork substrate shown in Fig. 1A. In these events, DNA extension temporarily exceeds the initial level before a final decrease. Dotted lines indicate the initial extension level.

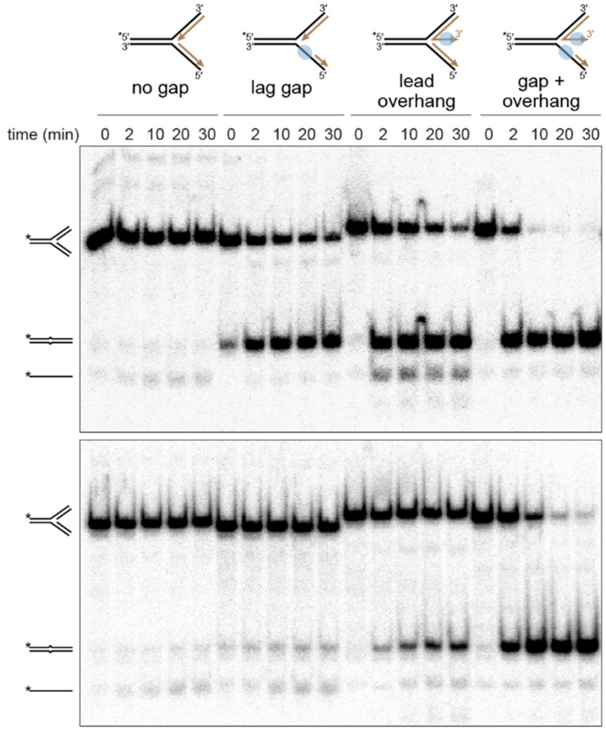

**Fig. S5. Fork reversal by SCF$^{FBH1}$ is stimulated by binding to 3'-ssDNA on the nascent leading strand.** Representative native polyacrylamide gel of fork reversal reactions of wild-type (top) or R447A K448A (bottom) SCF$^{FBH1}$ on fork substrates containing ssDNA binding sites on either the lagging strand template and/or the 3'-end of the nascent leading strand. Data from three independent experiments are quantified in Fig. 1F.

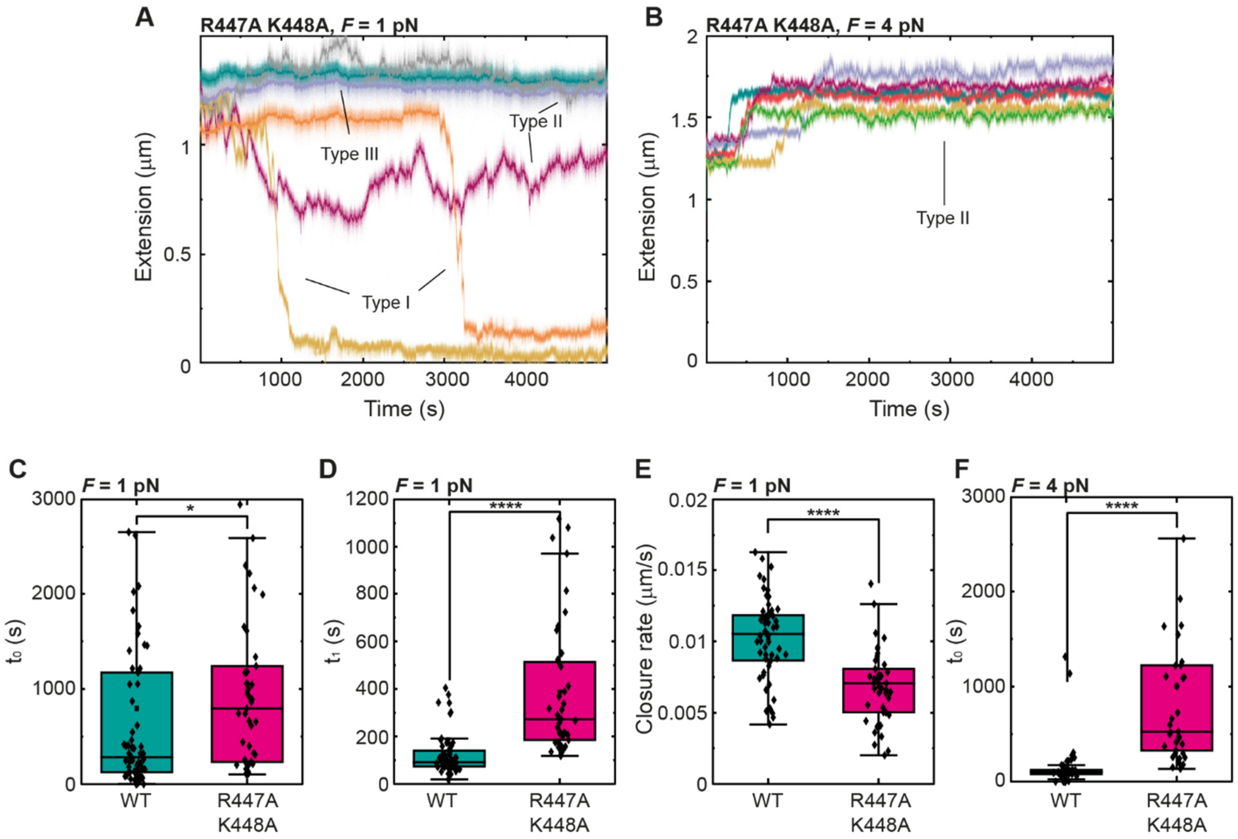

**Fig. S6. The R447A K448A mutation reduces SCF$^{FBH1}$ efficiency in fork reversal. A,B**. Representative time-extension traces showing individual activities of the SCF$^{FBH1}$ R447A K448A mutant (40 nM) on the fork substrate shown in Fig. 1A under 1 pN (A) and 4 pN (B) opposing force. **C.** Initial activity detection times ($t_0$) for wild-type (WT) SCF$^{FBH1}$ and the R447A K448A mutant at 1 pN force. p = 0.01405 as defined by Mann Whitney Test. **D.** Reaction completion times from the onset of the final extension decrease ($t_1$) for both proteins at 1 pN. p = 8.6E-12 as defined by Mann Whitney Test **E.** Closure rates of the fork substrate at 1 pN force. p= 9.7E-8 as defined by Two-sample t test. **F.** Initial activity detection times ($t_0$) at 4 pN force. p = 4.3E-14 as defined by Mann Whitney Test.

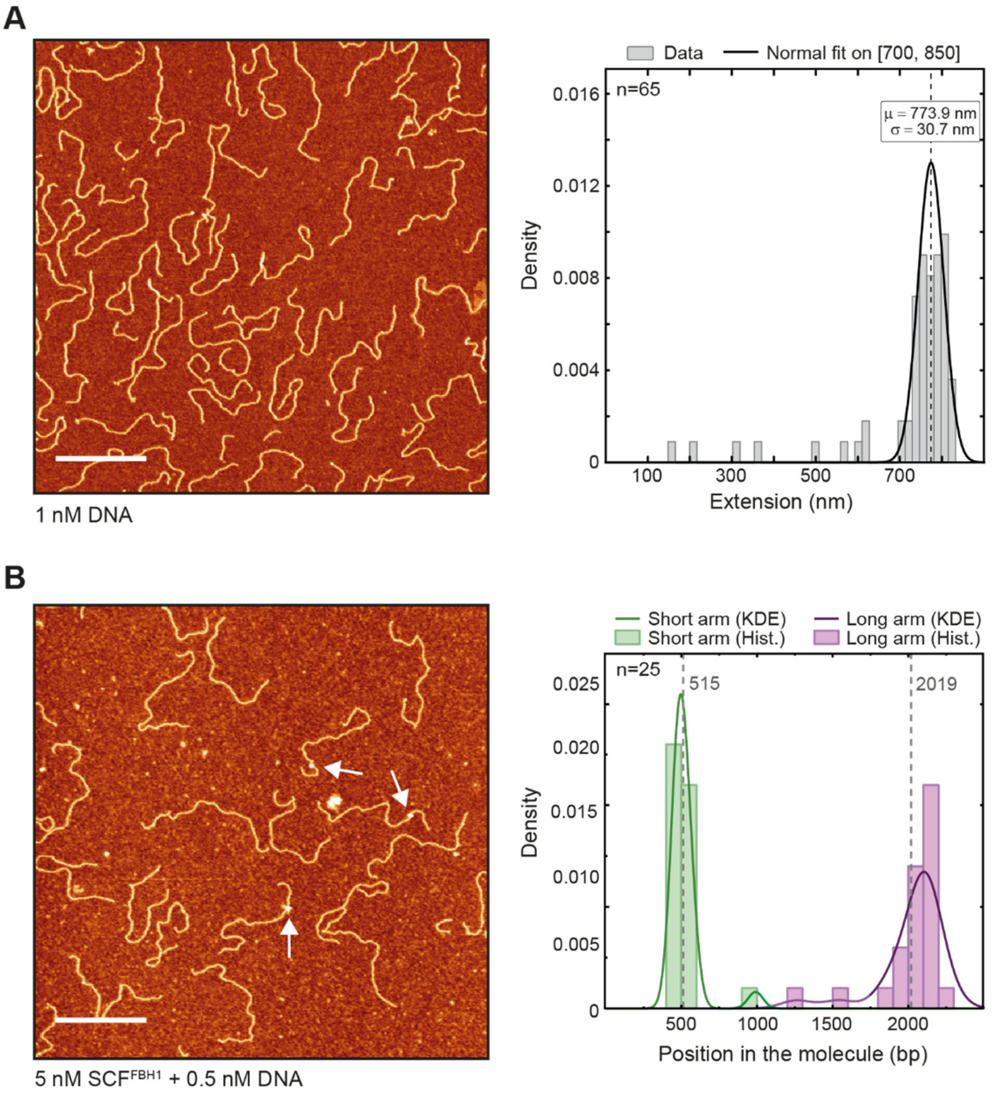

**Fig. S7. Controls for fork-reversal imaging. A**,**B**. AFM images (*left*) and distributions of dsDNA contour lengths (*right*) for the DNA fork substrate alone (A) or incubated with $SCF^{FBH1}$ (B). **A.** 1 nM DNA. A Gaussian fit applied over the 700–850 nm interval was used to determine the DNA contour length (dashed line). **B.** 5 nM $SCF^{FBH1}$ + 0.5 nM DNA. White arrows indicate molecules in which $SCF^{FBH1}$ interacts with the DNA. The plot quantifies the lengths of DNA arms emerging from the $SCF^{FBH1}$-bound junction in ATP-free reactions. Histograms show measured values; solid curves represent kernel density estimates (KDE). Dotted lines indicate the predicted arm lengths.

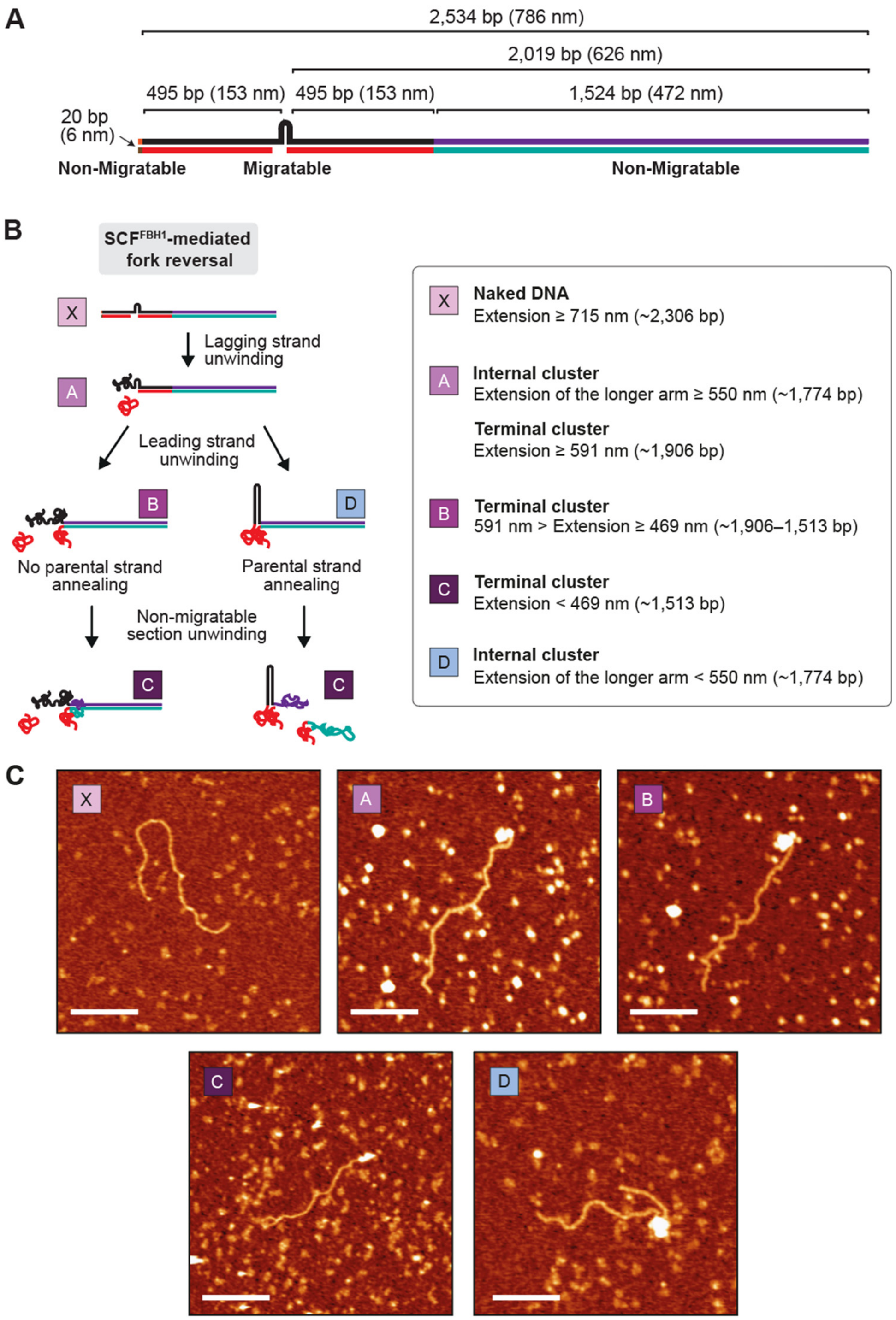

**Fig. S8. Reaction scheme and classification of SCF$^{FBH1}$-mediated reversal intermediates in the AFM substrate. A**. Schematic representation of the AFM DNA substrate. **B**. Classification criteria used to assign molecules for analysis of reaction progression, together with the corresponding schematic pathway of SCF$^{FBH1}$-mediated fork reversal on this substrate. **C**. Representative AFM images of each classified molecular species. Scale bar, 200 nm.

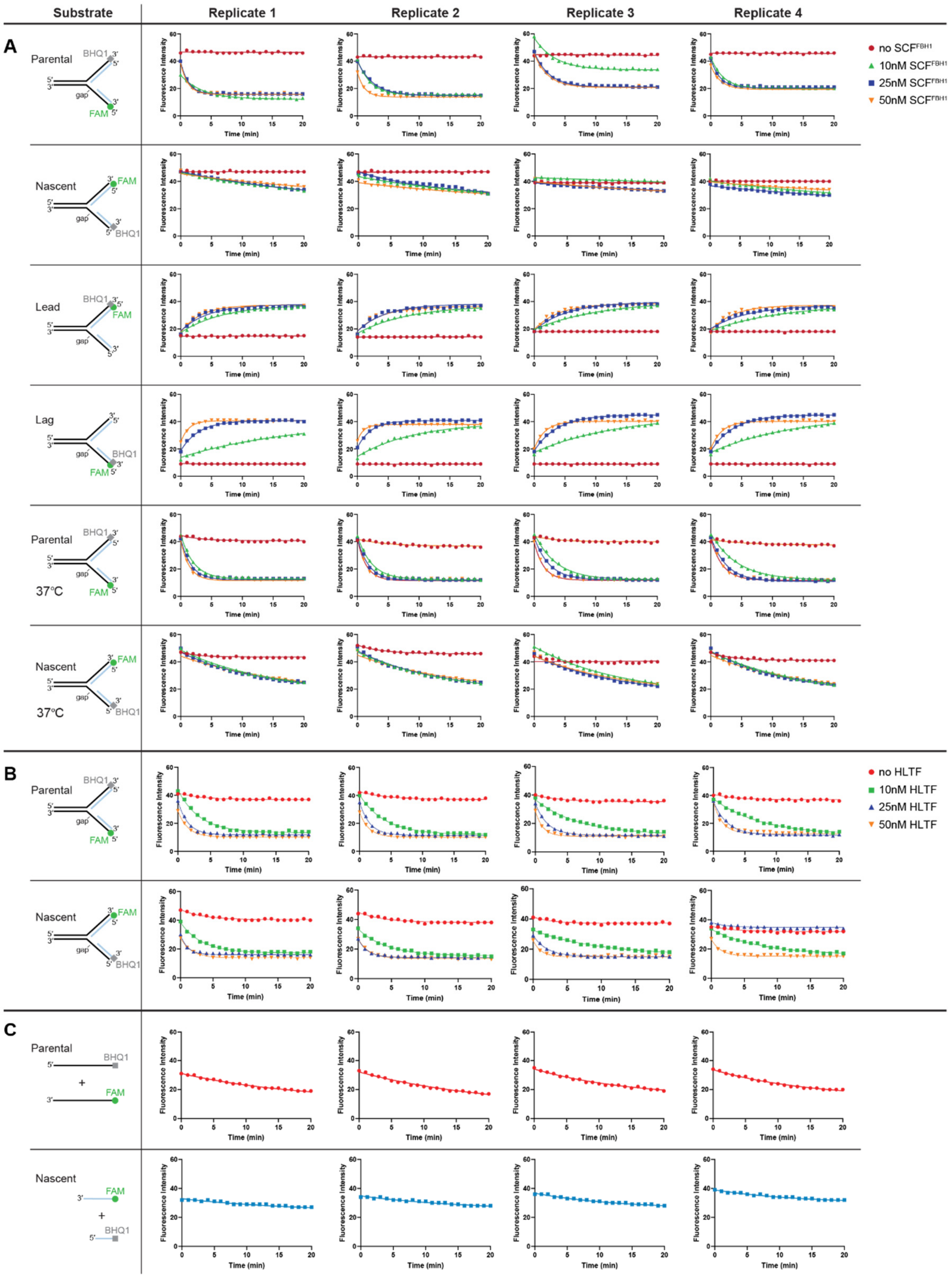

**Figure S9. Fluorescence traces for strand annealing and unwinding during fork reversal.** Four biological replicates are shown for each condition. **A,B**. Time-course fluorescence intensities for indicated substrates in the presence of varying concentrations of $SCF^{FBH1}$ (A) or HLTF (B). **C**. Control strand annealing reactions containing parental (top) or nascent (bottom) strands alone.

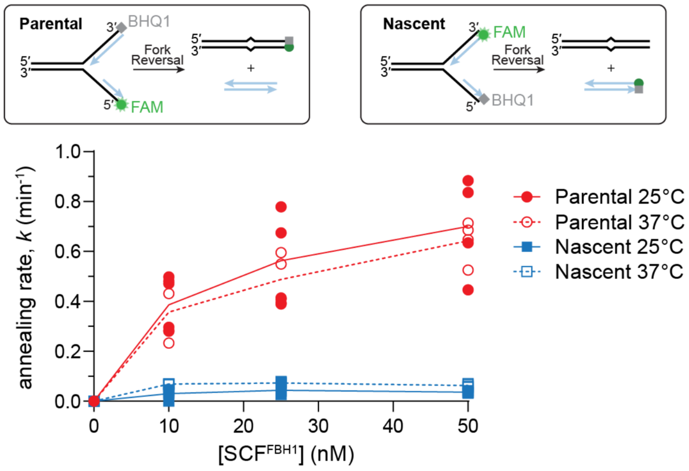

**Figure S10. SCF$^{FBH1}$ annealing rates are consistent at 25°C and 37°C.** Data at 25 nM SCF$^{FBH1}$ and 25°C are also shown in Figures 3C and 3D.

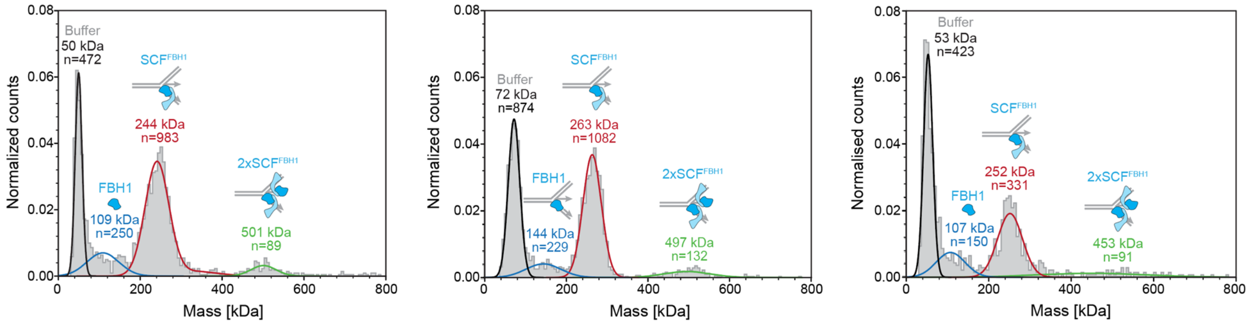

**Figure S11. Distribution of SCF$^{FBH1}$-fork complexes.** Mass photometry data for SCF$^{FBH1}$ in the presence of the *gap+overhang* substrate (three repeats). Histograms (grey) correspond to the normalized counts for each species present. Gaussian curves (black, blue, red, green) reflect the average mass of raw counts (n) for each peak, with colors corresponding to buffer (black), FBH1 (blue), 1×SCF$^{FBH1}$ bound to fork DNA (red), and 2×SCF$^{FBH1}$ bound to fork DNA (green). The FBH1 peak corresponded to either free FBH1 (left and right panels) or FBH1 bound to fork DNA (middle panel). Theoretical masses are as follows: fork DNA, 34 kDa; FBH1, 107 kDa; SCF$^{FBH1}$, 228 kDa.

**Supplementary Table S1. Oligodeoxyribonucleotides used in bulk biochemistry and structural studies**

| | | |
|---|---|---|
| **Fork Reversal[1]** | | |
| 1 | 48 | ($P^{32}$)ACGCTGCCGAATTCTACCAGTGCCTTGCTAGGACATCTTTGCCCACCTGCAGGTTCACCC |
| 2 | 50 | GGGTGAACCTGCAGGTGGGCAAAGATGTCC |
| 3 | 50_10polyT | GGGTGAACCTGCAGGTGGGCAAAGATGTCCTTTTTTTTTT |
| 4 | 52 | GGGTGAACCTGCAGGTGGGCAAAGATGTCCCAGCAAGGCACTGGTAGAATTCGGCAGCGTC |
| 5 | 53 | GGACATCTTTGCCCACCTGCAGGTTCACCC |
| 6 | 53_5gap | TCTTTGCCCACCTGCAGGTTCACCC |
| 7 | 48_Q_short | ATTCTACCAGTGCCTTGCTAGGACATCTTTGCCCACCTGCAGGTTCACCG(BHQ1) |
| 8 | 50_mod | CGGTGAACCTGCAGGTGGGCAAAGATGTCC |
| 9 | 52_F_short | (FAM)CGGTGAACCTGCAGGTGGGCAAAGATGTCCCAGCAAGGCACTGGTAGAAT |
| 10 | 53_5gap_mod | TCTTTGCCCACCTGCAGGTTCACCG |
| 11 | 48_short | ATTCTACCAGTGCCTTGCTAGGACATCTTTGCCCACCTGCAGGTTCACCG |
| 12 | 50_F | (FAM)CGGTGAACCTGCAGGTGGGCAAAGATGTCC |
| 13 | 52_short | CGGTGAACCTGCAGGTGGGCAAAGATGTCCCAGCAAGGCACTGGTAGAAT |
| 14 | 53_5gap_Q | TCTTTGCCCACCTGCAGGTTCACCG(BHQ1) |
| **EMSA** | | |
| 15 | FAM40 | (FAM)CTCAGGACTCAGTTCGTCAGCCCTTGACAGCGATGGAAGC |
| 16 | F20.40 | CGAAGGTAGCGACAGTTCCCCTGACGAACTGAGTCCTGAG |
| 17 | FAM40_lead2gap | GCTTCCATCGCTCTCAAG |
| 18 | FAM40_lag2gap | GAACTCTCGCTACCTTCG |
| 19 | FAM40_lag8gap | TCGCTACCTTCG |
| 20 | FAM40_lead2gap_10T | GCTTCCATCGCTCTCAAGTTTTTTTTTT |
| **Mass Photometry and EM** | | |
| 21 | FBH1_parental_leadFAM | (FAM)CTCAGTTCGTCAG CCCTAGACAGCGATG |
| 22 | lead_2gap_15tail | CATCGCTGTCTAGTCCACTCGTATTTGC |
| 23 | FBH1_parental_lag | GCGAAGGTAGCGACAGATCCCCTGACGAACTGAG |
| 24 | FBH1_lag_8gap_5T | TTTTTTCGCTACCTTCGC |
| 25 | FBH1_parental_lead | CTCAGTTCGTCAGCCCTAGACAGCGATG |

[1] Underlined nucleotides denote mismatches designed to prevent spontaneous fork reversal.

**Supplementary Table S2. DNA substrate construction**

| Figure | Substrate name | Annealed oligos[1] |
|---|---|---|
| **1F** | No gap | 1+2+4+5 |
| **1F** | Lag gap | 1+2+4+6 |
| **1F** | Lead overhang | 1+3+4+5 |
| **1F** | Gap+overhang | 1+3+4+6 |
| **3A-D** | Parental | 7+8+9+10 |
| **3A-D** | Nascent | 11+12+13+14 |
| **3E-G** | Lead | 7+10+12+13 |
| **3E-G** | Lag | 8+9+11+14 |
| **4A** | No gap | 15+16+17+18 |
| **4A** | Lag gap | 15+16+17+19 |
| **4A** | Lead overhang | 15+16+18+20 |
| **4A** | Gap+overhang | 15+16+19+20 |
| **4B** | Gap+OH Fork | 21+22+23+24 |
| **4C** | Gap+OH Fork | 22+23+24+25 |

[1] Oligo numbers from Table S1.

**Supplementary Table S3. Oligodeoxyribonucleotides used in single-molecule studies**

| Fragment | Oligonucleotide | Sequence |
|---|---|---|
| **MT Fork substrate** | | |
| PCR of dsDNA fragment connected to DIG handle | 401-230 HPLC | GGCGGCGAGCTGAGGGTTACCGGATAAGGCGCAGCG |
| | 381.R JY0 137 PspOMI-KpnI | ACTTACGCGGGCCCATCGACCGGCAATGAAGCAATGTTGCCAGCATTATGCAGGCCTGG |
| PCR of dsDNA fragment connected to BIO handle | 435.F ori 8 nts-BbvCI | TGACAAGAGGCGGCGAGCTGAGGGTTACCGGATAAGGCGCAGCG |
| | 296.R JY0 137 PspOMI-KpnI | ACTTACGCGGGCCCGGTACCGCCAGCATTATGCAGGCCTGGA |
| DIG side adaptor | 434.P-hairpin 8nt ov 137A | [Pho]TCAGCTCGCCGCCTCTTGTCACAGCAAGGCACTGGTAG |
| BIO side adaptor | 436.P-hairpin over 137Bv8 | [Pho]CCGACTACCAGTGCCTTGCTATGACAAGAGGCGGCGAGC |
| Short dsDNA hairpin | 250.Loop hairpin | [Pho]TCGGGTCAGATGCCTTTTGGCATCTGAC |
| MT Fork DIG handle | 57.FMH_F2_BamHI-ApaI | GCGTAAGTGGATCCGGGCCCCGACTCACTATAGGGAGACCGGC |
| | JOE-R1 | AGTAAGCGCCGTCAGACCAG |
| MT Fork BIO handle | 42.FMH_F2_KpnI-PsiI-ScaI | GCGTAAGTGGTACCTTATAAAGTACTCGACTCACTATAGGGAGACCGGC |
| | 209.BsrGI 71short handle | CGATAACCAACTGGCGATG |
| **Gapped substrate** | | |
| Complementary to released BbvCI fragment 1 | 137.block piece1 BbvCI | GGATGACATGAGCTGA |
| Complementary to released BbvCI fragment 2 | 138.block piece2 BbvCI | GGGTCAAGTGTGCTGA |
| Complementary to released BbvCI fragment 3 | 139.block piece3 BbvCI | GGCTAGCTGAGCTGA |
| Complementary to released BbvCI fragment 4 | 140.block piece4 BbvCI | GGTGATTGTAGGCTGA |
| MT gapped DIG handle | 57.FMH_F2_BamHI-ApaI | GCGTAAGTGGATCCGGGCCCCGACTCACTATAGGGAGACCGGC |
| | JOE-R1 | AGTAAGCGCCGTCAGACCAG |
| MT gapped BIO handle | FMH_F2_BsrGI | GCGTAAGTTGTACACGACTCACTATAGGGAGACCGGC |
| | 209.BsrGI 71short handle | CGATAACCAACTGGCGATG |
| **AFM Fork substrate** | | |
| PCR of dsDNA fragment connected to DIG handle | 401-230 HPLC | GGCGGCGAGCTGAGGGTTACCGGATAAGGCGCAGCG |
| | 403.R JY0 137s PspOMI-KpnI | ACTTACGCGGGCCCGGTACCCTCGCTCACTGACTCGCTGCG |
| PCR of dsDNA fragment connected to BIO handle | 435.F ori 8 nts-BbvCI | TGACAAGAGGCGGCGAGCTGAGGGTTACCGGATAAGGCGCAGCG |
| | 296.R JY0 137 PspOMI-KpnI | ACTTACGCGGGCCCGGTACCGCCAGCATTATGCAGGCCTGGA |
| DIG side adaptor | 434.P-hairpin 8nt ov 137A | [Pho]TCAGCTCGCCGCCTCTTGTCACAGCAAGGCACTGGTAG |
| BIO side adaptor | 436.P-hairpin over 137Bv8 | [Pho]CCGACTACCAGTGCCTTGCTATGACAAGAGGCGGCGAGC |
| Short dsDNA hairpin | 250.Loop hairpin | [Pho]TCGGGTCAGATGCCTTTTGGCATCTGAC |
